# Nucleo-cytoplasmic environment modulates spatio-temporal p53 phase separation

**DOI:** 10.1101/2023.10.16.562512

**Authors:** Debalina Datta, Ambuja Navalkar, Arunima Sakunthala, Ajoy Paul, Komal Patel, Shalaka Masurkar, Laxmikant Gadhe, Shinjinee Sengupta, Manisha Poudyal, Jyoti Devi, Ajay Singh Sawner, Pradeep Kadu, Ranjit Shaw, Satyaprakash Pandey, Semanti Mukherjee, Nitisha Gahlot, Kundan Sengupta, Samir K Maji

**Affiliations:** Department of Biosciences and Bioengineering, IIT Bombay, Powai, Mumbai-400076, India; Sunita Sanghi Centre for Aging and Neurodegenerative Diseases, IIT Bombay, Powai, Mumbai 400076; Amity Institute of Molecular Medicine & Stem Cell Research, AUUP, Noida, India; Chromosome Biology Lab, Indian Institute of Science Education and Research, Pune, India

**Author notes:** Correspondence: Prof. Samir K. Maji, Department of Biosciences and Bioengineering, IIT Bombay, Powai, Mumbai 400 076, India.

## Abstract

Phase separation of various transcription factors and nucleic acids into biomolecular condensates is known to play an essential role in the regulation of gene expression. Here, we show that p53, a tumor suppressor and transcription factor, phase separates and forms biomolecular condensates in the nucleus of cancer cells as well as when overexpressed in the various cell lines. Although the nuclear condensates of wild-type (WT) p53 maintain their liquid state and are able to bind DNA, cancer-associated mutations not only promote misfolding but also partially rigidify the p53 condensates, which are unable to bind the DNA. Irrespective of WT or mutant form, the cytoplasmic partitioning of p53 with time also results in biomolecular condensate formation, which eventually undergoes rigidification. *In vitro*, WT p53 core domain (p53C) forms biomolecular condensates, which rigidify with time and the process is further promoted by cancer-associated mutations. Both RNA and non-specific DNA promote LLPS of p53C, but specific DNA promotes the dissolution of p53C condensates. The result suggests that the cellular microenvironment regulates p53 LLPS, material property and its functions.

## Introduction

The cell is organized into sub-cellular compartments for carrying out specific biological reactions, which is essential for the cell’s functions and survival^1,2^. While several of these compartments are membrane-bound, multiple membraneless organelles exist, including nucleoli, PML bodies and centrioles^3–6^. Studies over the last decade have revealed that liquid-liquid phase separation (LLPS) leading to biomolecular condensate formation is a critical phenomenon involved in the formation, function, and regulation of these membraneless organelles in the cell^5–13^. Some well-known examples of biomolecular condensates are nuclear speckles, stress granules and cell signaling clusters, which play various essential roles in cellular activities, including transcriptional regulation, cellular stress response and signal transduction^4,14–19^. Proteins containing intrinsically disordered regions (IDRs) or prion-like domains, in the presence or absence of other biomolecules such as DNA/RNA, are known to undergo phase separation and form biomolecular condensates^12,20–28^. Apart from having physiological functions, LLPS is also known to be associated with disease pathology spanning from neurodegenerative diseases to cancer^5,29–31^.

Previously, it has been shown that many transcription factors undergo LLPS, where a phase-separated condensate state is linked to their transcriptional activities^4,32,33^. p53 is one such transcription factor, which regulates many biological processes, including cell cycle, DNA repair and apoptosis^34–36^. Moreover, the transcriptional regulator p53 plays an essential role in tumor suppression^37–41^. This is further supported by the fact that most cancers are often characterized by p53 mutations wherein p53 frequently accumulates in the nucleus as well as in the cytoplasm^42–46^. Recent studies suggested that p53 misfolding, aggregation and amyloid formation, which are further promoted by cancer-associated mutations, are shown to be associated with p53 loss of function and gain of oncogenic functions^45,47–52^. Further, it was shown that p53 phase separation into biomolecular and/or mesoscopic condensates could act as a potential precursor for p53 amyloid formation^53–55^. Moreover, the partitioning of p53 into cellular condensates such as Cajal and promyelocytic leukemia protein (PML) bodies may also regulate the post-translational modifications and function of p53^56,57^.

p53 is a nuclear phosphoprotein, and its nuclear localization is essential for growth-suppressing activity in the late G1 stage^58,59^ of the cell cycle. In normal cells, p53 levels are tightly regulated as a result of a short half-life (15 to 30 min)^60^ and p53 is undetectable by immunocytochemistry^61^. Upon stress signals such as DNA damage, p53 gets stabilized and its local concentration increases in the nucleus for its tumor suppressive functions^62^. We hypothesized that p53, upon its stabilization and/or high expression, may undergo LLPS and form biomolecular condensate in the nucleus, which might act as a local reservoir of the functional protein for immediate functions such as DNA binding. Indeed, the STED microscopy of two cancer cells (MDA-MB-231 and MCF7 cells) with known nuclear p53 stabilization as well as overexpression of p53 in HeLa and SaOS2 cells (p53 null) showed p53 condensate formation in the nucleus. The overexpression cellular model suggests initial p53 condensates formation predominantly localized in the nucleus in a liquid-like state, however, with time, cytoplasmic p53 condensates appear, which transition into an arrested state. The spatiotemporal formation, modulation (cisplatin treatment) and material property of p53 condensate may modulate p53 function and cell fate. In fact, the oncogenic mutants p53 R175H and p53 R248Q show an enhanced transition to a dynamically arrested state both in the nucleus and the cytoplasm, which inhibits their binding to p53 specific DNA. Consistent with cellular data, the p53 core domain (DNA-binding domain, p53C) also undergoes LLPS *in vitro* and shows rigidification with time, which is further promoted by cancer-associated p53 mutations. Based on the observation that nuclear p53 condensates maintain the liquid-like state over time, whereas cytoplasmic condensates undergo transition to an arrested state, we hypothesized that the nucleo-cytoplasmic microenvironment (particularly DNA/RNA) might modulate the formation and material properties of p53 condensates. Indeed, the presence of both RNA and non-target DNA (NTR-DNA) promotes the p53C phase separation and maintains liquid-like properties, whereas the target DNA (TR-DNA) promotes the soluble state of p53. Overall, our study provides mechanistic insights into the physical state of the p53 condensates in the cell, its correlation to p53 transcriptional function and cell fate.

## Results

### p53 forms liquid condensates in cancer cells

Tumor suppressor, p53, is a 393-amino acid protein and transcriptional factor consisting of four major domains: transactivation domain (TAD), proline-rich region (PRR), DNA-binding domain (DBD) and tetramerization domain (TD)^63^ **(Fig. 1a)**. Bioinformatics analysis suggests that p53 not only contains low complexity region (63-89 residues) but also possess multiple disordered regions (residues 30-98, 283-328 and 343-393) **(Fig. 1a)**. The droplet forming propensity of p53 predicted by FuzDrop^64^ revealed four major droplet-promoting regions at the N-terminus (residues 1-24 and 28-108) and C-terminus (amino acids 227-337 and 341-393) **(Fig. 1a)**. Interestingly, the secondary structure prediction by Alphafold^65^ and recent experimental studies^66^ revealed that the N and C-terminal of the protein majorly comprise random coil structure, whereas the core DNA-binding region consists of α-helix and β-sheet conformation **(Fig. 1b)**.

**Figure 1.**
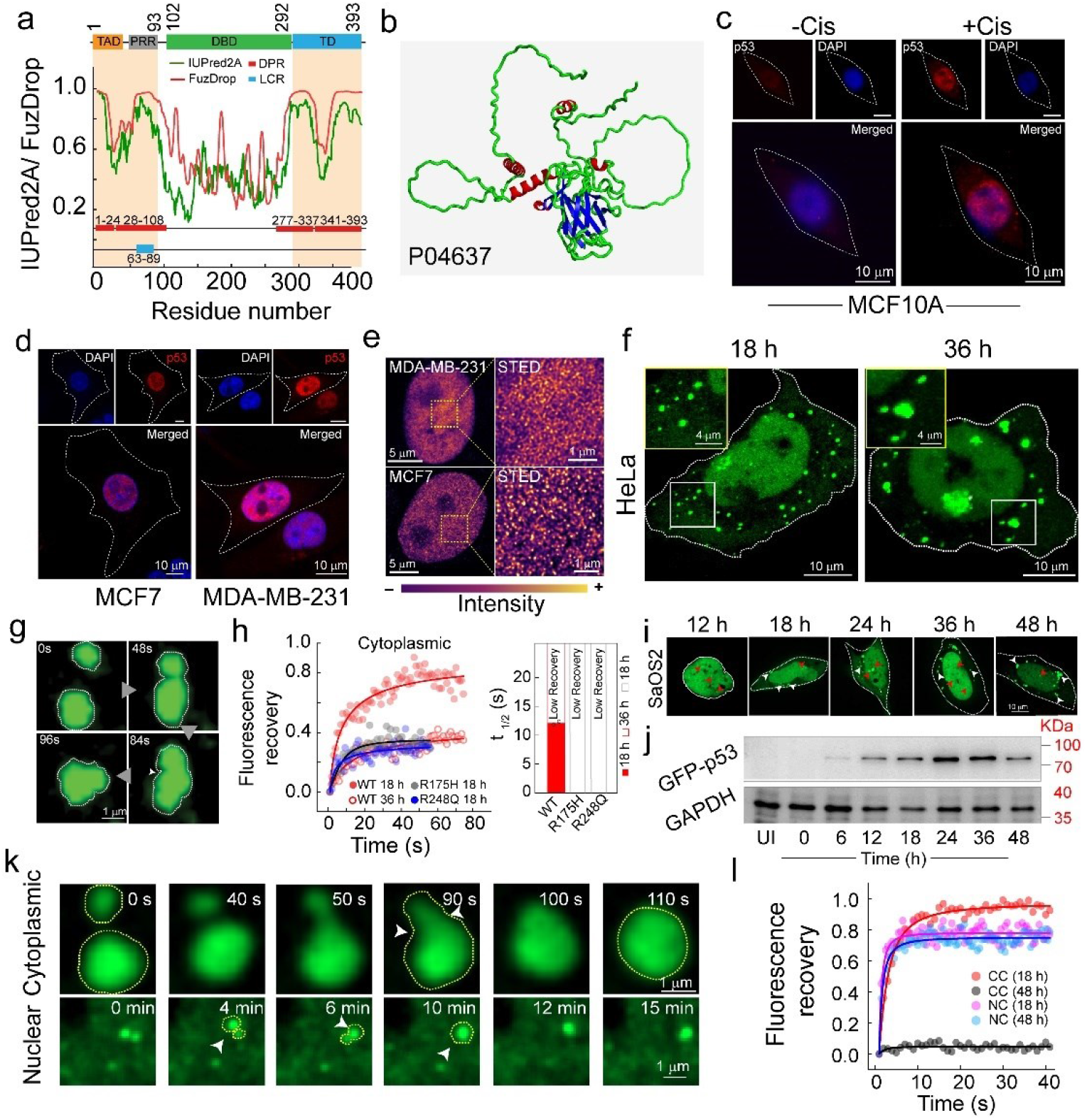
p53 condensate formation in HeLa and SaOS2 cells. **(a)** Bioinformatics analysis to predict the phase separation behaviour of p53. **(*Upper panel*)**, Domain organization of p53 showing TAD (1-61), PRR (64-93), DBD (94-292) and TD (325-393). (***lower panel***) *In silico* analysis of the primary sequence of p53 showing the presence of low complexity regions (blue) using SMART, disorder tendency (green) using IUPRED2A, residue-based droplet promoting probabilities (red) and droplet-promoting regions (DPR) (red box) using FuzDrop. **(b)** Structure of full-length p53 protein as predicted by Alphafold from 229 structures in PDB (Uniprot ID P04637), showing the organization of its secondary structure (red, green, and blue colour represents α-helix, random coil and β-sheet conformation, respectively). **(c)** Representative confocal images of MCF 10A cells stained with p53 antibody (DO-1) showing stabilization of p53 (red) in the nucleus after cisplatin treatment for 3 h ***(right panel)*** compared to untreated cells ***(left panel)***. The scale bar is 10 μm. **(d)** Immunostaining using p53 antibody (DO-1) followed by confocal microscopy showing the localization of p53 (red) in the nucleus in breast cancer cell lines MCF7 and MDA-MB-231. The scale bar is 10 μm. **(e)** Super-resolution imaging using STED microscopy of MDA-MB-231 and MCF7 cells after immunostaining with p53 antibody (DO-1) shows p53 in the condensate state throughout the nucleus in both cell lines. Pseudo color (LUT, mpl-plasma) has been used for representative purposes. The scale bar represents 1 μm. **(f)** Representative confocal images of HeLa cells over-expressing WT GFP-p53 at 18 h and 36 h showing the presence of cytoplasmic condensates. The zoomed inset (18 h) shows the intracellular spherical condensates at early time point and enhanced accumulation of condensates with loss of spherical structure at later time point. The scale bar is 10 μm. The corresponding video for 18 h is provided as Supplementary Movie 1. **(g)** Time-lapse microscopic images showing fusion of WT GFP-p53 condensates in the cytoplasm of HeLa cells captured at 18 h post-transfection. The scale bar is 1 μm. The corresponding time-lapse video for fusion is provided as Supplementary Movie 2. **(h)** Fluorescence recovery after photobleaching (FRAP) curves of cytoplasmic condensates in HeLa cells at 18 h and 36 h showing fluorescence recovery with ***(right panel)*** their corresponding t_1/2_. Note that the t_1/2_ for condensates with low recovery was not estimated. **(i)** Time-dependent expression of WT GFP-p53 in SaOS2 cells, showing the temporal evolution of cytoplasmic condensates in SaOS2 cells where distinct nuclear condensates (red arrows) and cytoplasmic condensates (white arrows) were seen. The scale bar is 10 μm. **(j)** Western blot images showing the expression of GFP-p53 with time in SaOS2 cells. **(k)** Time-dependent confocal images of cytoplasmic and nuclear condensates of WT GFP-p53 showing the fusion event of two condensates. The scale bar is 1 μm. The corresponding video for the fusion of cytoplasmic condensates is provided as Supplementary Movie 3. **(l)** FRAP curves of both nuclear (NC) and cytoplasmic condensate (CC) showing almost 100% recovery (18 h) and no recovery (48 h) for cytoplasmic condensates, whereas nuclear condensates showed complete recovery at both 18 h and 48 h. All the experiments **(c-l)** were repeated two times with similar observations.

In healthy cells, the basal level of p53 remains low due to its rapid turnover^67^. In the presence of stress (such as DNA damage), p53 undergoes post-translational modifications^68,69^ and is stabilized^70^, which is eventually transported to the nucleus for the functions. In fact, when epithelial cells MCF-10A is treated with cisplatin (DNA damage agent^71^), we observed nuclear p53 stabilization in contrast to untreated cells **(Fig. 1c)** using confocal microscopy. This is also consistent with p53 status in two cancer cells (MDA-MB-231 and MCF7 cells) **(Fig. 1d)**, where nuclear stabilization of functional p53 is known^47^. To further examine the nuclear p53 state, we performed super-resolution imaging studies using stimulated emission depletion microscopy (STED)^72,73^ with MDA-MB-231 and MCF7 cells **(Fig. 1e)**. The STED microscopy revealed the presence of dense nuclear p53 condensate-like state **(Fig. 1e)**. This indicates that p53 might naturally exist in phase separated state when stabilized or overexpressed in cells. These phase separated p53 in the nucleus are unlikely to be amyloidogenic in nature as nuclear p53 in these cells was previously shown to be OC (amyloid specific antibody) negative^47^. Further, when GFP-p53 WT is overexpressed in HeLa cells, p53 showed majorly nuclear localization (∼12 h) (**Fig. S1b**), however at later time points (∼18 h onwards), the cells showed cytoplasmic condensates (**Fig. 1f, Fig. S1a, b and Supplementary Movie 1**). At this stage, we only observed distinct p53 condensate formation in the cytoplasm, however intense p53 signal with few p53 condensates were visible in the nucleus. HeLa cells expressing only GFP did not show any appearance of condensates but rather showed a diffuse pan-cellular signal confirming that liquid condensate formation is a property of p53 and not influenced by the GFP tag (**Fig. S1c**).

Since cytoplasmic condensates were distinct, further characterizing of these condensates showed that at the early time points (18 h), they were highly dynamic, spherical in nature (diameter ∼1 µm) and exhibited fusion to form large assemblies (diameter ∼3 µm) **(Fig. 1g and Supplementary Movie 2)** suggesting their liquid-like behavior. This is further supported by their high fluorescence recovery after photobleaching (FRAP) (∼ 80 % recovery) with a short t_1/2_ (∼12 s) (**Fig. 1h and Fig. S1d**). At a later time point (36 h), p53 condensates in the cytoplasm not only lose their spherical shape (**Fig. S1e**) but also possess a substantial decrease in FRAP recovery (∼30%), suggesting their transition towards an arrested state. Interestingly, once overexpressed, condensates formed by two cancer-associated hotspot p53 mutants (R175H and R248Q) exhibited an arrested state (very low FRAP recovery) at the early time point (18 h) **(Fig. 1h and Fig. S1d, f)**. To examine the protein expression level, Western blot analysis was done, which showed a gradual increase in expression of GFP-p53 (up to 36 h) that was sustained till 48 h **(Fig. S1g, h)**. The data, therefore, indicates that upon over expression, p53 is retained in the nucleus, but with time, p53 partitions into the cytoplasm, where it undergoes liquid-to-arrested state transition **(Fig. S1a and b)**.

To nullify the effect of endogenous p53 and to establish functional consequences due to p53 condensate formation, we examined p53 LLPS in p53 null SaOS2 cells^74,75^. Similar to HeLa cells, we observed that at an early point (12 h), p53 was majorly localized in the nucleus with the appearance of a few p53 condensates. With time, the distinct cytoplasmic condensates of GFP-p53 started to appear after 18 h of transfection (**Fig. 1i**). Important to note that SaOS2 cells expressing only GFP did not show any condensate formation (**Fig. S2a**) suggesting that GFP do not influence the p53 condensate formation similar to HeLa cells. Along with the confocal microscopy, the Western blot analysis showed a sustained increase in the expression of p53 from 6 to 24 h, after which it slightly reduced until 48 h (**Fig. 1j and Fig. S2b)**. The early formed p53 condensates were liquid-like (for both nuclear and cytoplasmic condensates) as confirmed by their fusion events **(Fig. 1k and Supplementary Movie 3)** as well as high FRAP recovery (∼80-100%) (**Fig. 1l**). Interestingly, FRAP studies at the late time point (48 h) showed, although nuclear condensates exhibit high recovery (∼80%), the cytoplasmic condensates showed almost no recovery (less than 10%) (**Fig. 1l and Fig. S2c, d**). These data suggest that once p53 is expelled out of the nucleus, the condensates undergo a liquid-to-arrested state transition in the cytoplasmic environment. This is further consistent with morphological analysis, where cytoplasmic condensates showed an increase in size and a decrease in circularity with time (**Fig. S2e**).

### Spatio-temporal partitioning of p53 condensates

To understand nucleo-cytoplasmic partitioning and condensate formation of p53 with time, we performed live-cell imaging experiments with SaOS2 cells. At the early time point, the p53 signal was mostly localized in the nucleus, but with time, p53 appeared in the cytoplasm and formed cytoplasmic condensates. (**Fig. 2a and Supplementary Movie 4**). The increased cytoplasmic/nuclear (C/N) ratio of the GFP-p53 signal with time indicates the possibility of nucleo-cytoplasmic shuttling of p53 (**Fig. 2b**). This is further confirmed by the reduction of C/N ratio on treatment with Leptomycin B (LMB, inhibitor of nuclear export^76^) indicating p53 transports to the cytoplasm through nuclear pore complex **(Fig. 2c)**. Indeed, GFP-p53 NES-overexpression in SaOS2 cells showed only nuclear localization of p53 (from 18 h to 48 h) (**Fig. 2d and Fig. S2f)**. These data suggested that the cytoplasmic p53 condensates are formed after expulsion of p53 from the nucleus. These cytoplasmic condensates did not exhibit any colocalization with Nile red (specific for intracellular lipid^77^), Lysotracker red (for lysosomes) and Mitotracker red (for mitochondria), suggesting their membraneless state **(Fig. S3)**. Moreover, they also do not remain associated with the aggresome formation in the cytoplasm **(Fig. S3)**.

**Figure 2.**
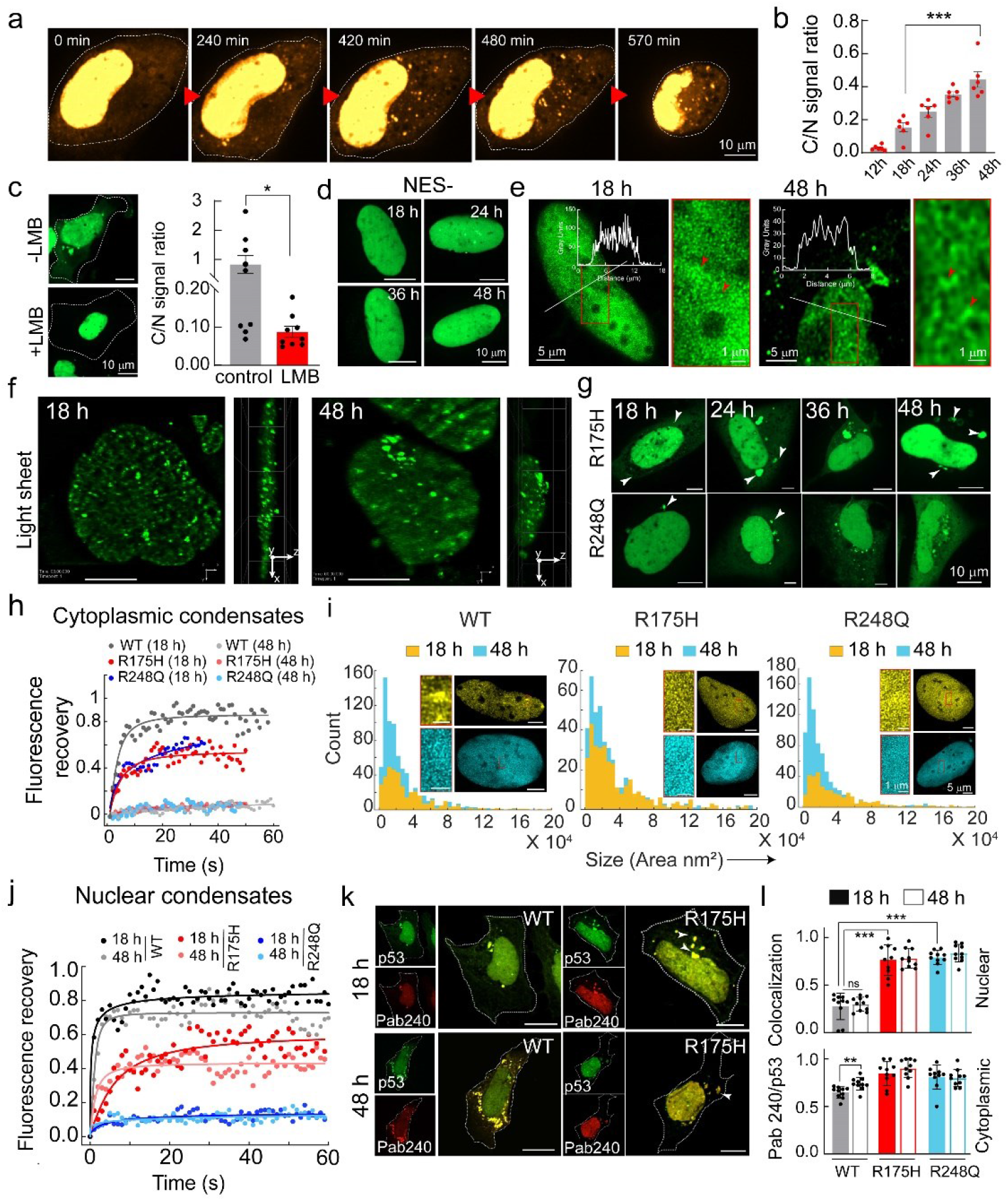
Spatio-temporal formation of p53 condensates in p53-null SaOS2 cells. **(a)** Time-lapse confocal microscopy images showing spatio-temporal localization and formation of p53 condensates. At early time points, only nuclear p53 signals were seen, followed by the appearance of cytoplasmic condensates and cell death. The scale bar is 10 μm. The corresponding time-lapse video is provided as Supplementary Movie 4. **(b)** Cytoplasmic/Nuclear (C/N) ratio of WT GFP-p53 fluorescence signal in SaOS2 cells with time (estimated based on intensity calculations) showing a gradual increase in GFP fluorescence signal in the cytoplasm. Error bars represent mean ± s.e.m for *n=6* transfected cells. **(c)** Leptomycin B (LMB) treatment of SaOS2 cells post-transfection with WT GFP-p53 showed reduced cytoplasmic p53 intensity compared to the untreated control. The scale bar is 10 μm. ***Left panel:*** Representative confocal microscopy images of treated cells and untreated control. ***Right panel***: The C/N ratio is estimated after image analysis. Error bars represent mean ± s.e.m for *n=9* transfected cells. **(d)** The time-lapse confocal microscopy showed only nuclear localization of the p53 signal after transfection with p53 NES-in SaOS2 cells. The scale bar is 10 μm. **(e)** Super-resolution microscopy (STED) of nuclear condensates of WT GFP-p53 showing the condensates of p53 in the nucleus (18 h and 48 h). The scale bar is 5 μm. The line profile represents the intensity of GFP-p53 across the nucleus. **(f)** Lattice light-sheet imaging of nuclear condensates of SaOS2 transfected with WT GFP-p53 at 18 h and 48 h. The scale bar represents 10 μm. The corresponding video showing the dynamic nature of the nuclear condensates is provided as Supplementary Movie 5. **(g)** p53 condensate formation by cancer-associated mutations of R175H and R248Q transfected in SaOS2 cells monitored over time using confocal microscopy. Scale bar is 10 μm. (**h**) FRAP recovery profiles of cytoplasmic condensates of WT, R175H and R248Q showing complete recovery of WT while low fluorescence recovery by mutants at 18 h and no recovery by all these proteins at 48 h. (**i**) Size (area) and number distribution of nuclear condensates of WT, R175H and R248Q calculated from STED microscopy showing an increase in the condensate number without altering the size at late (48 h) time point compared to early (18 h) time point by all these proteins. Scale bar is 5 μm. **(j)** FRAP recovery profile of nuclear condensates of WT, R175H and R248Q showing complete recovery of WT p53 condensate at early (18 h) and late time point (48 h) while partial recovery of R175H and no recovery for R248Q. **(k)** Immunofluorescence study showing the lesser extent of colocalization of WT GFP-p53 condensates with misfolded p53 specific antibody (Pab240), while higher colocalization of the same for R175H GFP-p53 condensates. The scale bar is 10 μm. **(l)** Bar graph representing the fraction of Pab240 colocalization with GFP-p53 nuclear and cytoplasmic condensates of WT, R175H and R248Q at early (18 h) and late (48 h) time points. The plot represents the mean with s.e.m, for n=2 independent experiments. For **(b)**, **(c)** and **(l),** the statistical significance was calculated using a two-tailed t-test (p<0.001(***), p<0.002(**), p<0.033 (*) and p>0.012 (ns), with 95% confidence interval. The experiments **(a, c-h, j and k)** were performed two independent times.

To further characterise the p53 condensates in the nucleus and cytoplasm, STED microscopy of GFP-p53 in SaOS2 cells was performed. The data showed that the nucleus of the cells is rather crowded with p53 condensates, which sometimes clump together to form local regions of high density (**Fig. 2e**). Further analysis also revealed that the majority of the nuclear p53 (>70%) exists as condensates (**Fig. S2g**), where morphology appeared to be an interconnected microphase separated condensates (percolation clusters) as reported previously^78–80^. The STED microscopy of cytoplasmic condensates suggests that both WT and mutant p53 exist in the form of small condensates **(Fig. S4)**. To examine whether most of the p53 remains in a condensate state in live cells, we also performed lattice light-sheet microscopy of SaOS2 cells expressing WT GFP-p53. The data showed largely dynamic p53 nuclear condensates throughout the nucleus **(Fig. 2f and Supplementary Movie 5)**.

We further studied the effect of two hotspot p53 mutants associated with cancer (R175H and R248Q) and the data showed that they formed slightly larger cytoplasmic condensate (18 h) as compared to WT protein (∼1 µm), but their decrease in circularity with time was similar to WT protein (**Fig. 2g and Fig. S2h, i**). FRAP analysis of the cytoplasmic condensates showed, in contrast to WT p53, only ∼50% recovery was observed for both the mutants even at an early time point (18 h), which further reduced to almost no recovery at 48 h **(Fig. 2h, Fig. S2j)**. The data suggests a faster liquid-to-arrested-state transition of the mutant p53 condensates as compared to WT p53 in the cytoplasm.

When R175H and R248Q nuclear condensates were characterized using STED microscopy, the data revealed that both mutants formed a dense population of condensates in the nucleus at 18 h, having an apparent higher density of accumulation at certain local regions **(Fig. S5)**. At a late time point (48 h), larger sized (in terms of area parameter) and more population of nuclear condensates were seen for both mutants, similar to WT protein (**Fig. 2i**). Similar to the nucleus, we also did not find any major morphological difference in the cytoplasmic condensates between WT and mutants by STED microscopy **(Fig. S4)**.

Interestingly, when FRAP analysis was performed, both mutant condensates showed significantly reduced recovery of ∼40% (for R175H) and ∼10% (R248Q), respectively, at both time points (18 h and 48 h). This indicates an arrested nature of nuclear condensates of mutants even at an early time point (18 h) (**Fig. 2j, Fig. S2k**). Interesting to note that p53 R175H condensates showed a relatively higher FRAP recovery compared to R248Q.

Thus, the data suggest that while cytoplasmic condensates transitioned from liquid to an arrested state, which is further promoted by both cancer-associated mutants, the nuclear WT p53 condensates, however, remained in a liquid-like state in SaOS2 cells. The rigidification of nuclear condensates of p53 mutants and cytoplasmic condensates in both WT and mutants in SaOS2 cells could be due to stronger assembly of p53 caused by misfolding and/or aggregation.

Indeed, immunofluorescence study using Pab240 antibody (specific to misfolded p53^81^) showed that mutant p53 condensates in the nucleus were more in misfolded conformation compared to WT. Additionally, the majority of the cytoplasmic condensates (both WT and mutants) were misfolded, where the mutants showed a higher colocalization of p53 with Pab240 compared to WT (**Fig. 2k, l and Fig. S6a**). However, the amyloid-specific OC antibody staining did not show any colocalization with GFP-p53 WT and mutants GFP-p53 R175H and GFP-p53 R248Q **(Fig. S6b)**. This suggests that although p53 remains in a misfolded conformation in the cytoplasm, it does not show the presence of amyloid conformation.

### The functional consequence of p53 phase separation in cells

To investigate the consequences of p53 nuclear condensate formation on DNA binding and transcriptional activity, we monitored the p53 downstream gene p21 by Western blot analysis. The data suggest a time-dependent increase in p21 expression up to 24 h and then plateaued (**Fig. 3a, b**), indicating the presence of functional p53 in the nucleus. This is further apparent as Annexin-PI assay followed by FACS analysis showed a gradual increase in apoptotic population of cells with time (**Fig. 3c and Fig. S7**). This increase in p21 expression and apoptosis might be due to the higher expression of p53 with time **(Fig. 1)**. We further examine p53 binding to its direct target, p21 specific DNA sequence, by ELISA and ChIP assay at 18 h of transfection for both WT and mutants. While WT p53 showed higher p53 DNA binding, both cancer-associated mutants (R175H and R248Q), however, showed a much lesser extent of DNA-binding activity **(Fig. 3d, e)**. This is consistent with the fact that mutant p53 condensates are composed of misfolded p53, which promotes condensate rigidification even at 18 h **(Fig. 2j, k)**, where p53 is most likely to be non-functional. To further examine the effect of DNA damage response to p53 condensate (p53 activation), SaOS2 cells were transfected with WT GFP-p53 and treated with cisplatin. After 18 h, the STED imaging indicated a higher number of p53 condensates in the nucleus for cisplatin treated cells as compared to untreated cells (**Fig. 3f-h and Fig. S8**). Interestingly, increased p53 condensate formation is also correlated with increased p53 stabilization **(Fig. 3i, j)** and p21 mRNA levels **(Fig. 3k)** in cells upon treatment with cisplatin in comparison to untreated cells. The data suggest that the p53 condensates might be functional in terms of DNA binding activity, which might be further enhanced when DNA damage occurs for tumor suppressive functions.

**Figure 3.**
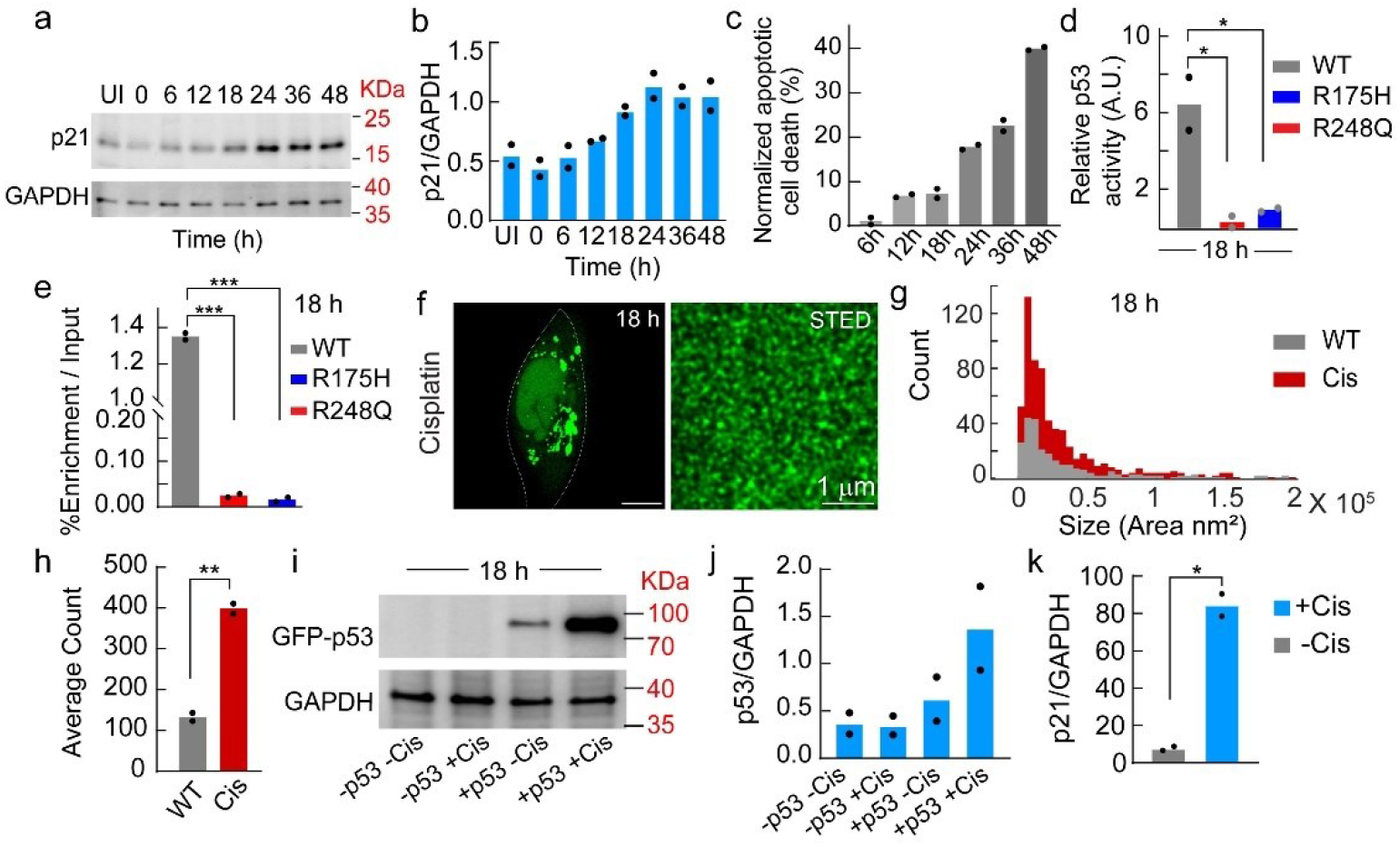
Stabilization and activity of p53 condensates. **(a)** Time-dependent expression of p21 (p53 target) using Western blot analysis showing a trend of p21 expression similar to p53. **(b)** Fold-change of p21 was quantified from the Western blot analysis. **(c)** Time-dependent apoptotic cell death of SaOS2 cells monitored by Annexin-PI assay followed by FACS analysis demonstrating an increase in cell death over time. **(d)** Evaluation of p53 DNA binding capacity of p21-specific DNA sequence by ELISA assay at 18 h for WT and p53 mutants. **(e)** Chromatin immunoprecipitation (ChIP) assay for evaluation of p53 DNA binding ability using p21-response element at 18 h showing higher DNA binding of WT p53 in comparison to p53 mutants. For ELISA and ChIP assays, statistical significance was determined by one-way ANOVA using Newman−Keuls (SNK) post hoc analysis (p<0.001(***), p<0.01(**), p<0.05 (*)). **(f)** SaOS2 cells transfected with WT GFP-p53 after treatment with cisplatin (stressor) showing p53 condensates at 18 h (***left panel***). Super-resolution STED imaging showing the presence of p53 clusters in the nucleus after treatment at 18 h (***right panel***). The scale bar is 1 μm. **(g)** Size/number distribution showing wide distributions of nuclear condensates for cisplatin-treated cells compared to untreated cells. **(h)** Estimation of condensates count in uniform area of (5μm x 5μm, n=2) condensates in the nucleus of cisplatin treated SaOS2 cells showing a higher number of condensates in treated (Cis) cells compared to untreated (WT) cells. **(i)** Western blot showing higher p53 expression after treatment with cisplatin (10 μM, 9 h) in comparison to untreated cells. **(j)** p53 levels were quantified from the Western blot images at the corresponding conditions. **(k)** The p21 mRNA expression of SaOS2 cells transfected with WT GFP-p53 at 18 h in the presence and absence of cisplatin treatment showing an increased level in the treated cells compared to the untreated control. For **(h)** and **(k),** the statistical significance was calculated using unpaired two-tailed t-test (p<0.001(***), p<0.002(**), p<0.033 (*) and p>0.012 (ns), and) with 95% confidence interval. The experiments **(a-f and i-k)** were repeated two independent times.

### p53 DNA-binding domain (p53 core, p53C) undergoes LLPS and liquid-to-arrested state transition *in vitro*

p53 core domain (p53C) represents the DNA binding domain of full-length p53, which is most often used for *in vitro* studies due to the unstable nature of full-length p53 protein ^82–84^. The previously reported crystal structure of p53C^85^ shows the organization of its secondary structure into α-helical, β-sheet, and random coil conformations (**Fig. 4a**). Further, this domain also contains intrinsically disordered regions (IDR, predicted by IUPRED 2A^86,87^) and two droplet-promoting regions (DPR, predicted by FuzDrop^64^), which are present at the N-terminus and C-terminus of the p53C (**Fig. 4b and Fig. S9a, b**). For *in vitro* phase separation, WT p53C was recombinantly expressed, purified (**Fig S9c, d**) and NHS-Rhodamine labelled p53C (10:90 labelled: unlabelled protein) was incubated in the presence of varying percentages of PEG-8000 (as molecular crowder) at pH 7.4 in 50 mM phosphate buffer. When examined under microscopy, we observed that both WT and mutants (R175H and R248Q) p53C readily phase separates to form condensates at as low as 10 μM concentration in the presence of 10% PEG-8000 (**Fig 4c, Fig. S9e**) with no significant difference in their phase regime **(Fig. 4d, Fig. S9f-h and Supplementary Movie 6)**. The mutant R175H showed a slightly higher propensity for phase separation as compared to WT and R248Q. The condensate formation was further examined by static light scattering (LS) experiments (at 350 nm), which showed all proteins spontaneously phase separate within a few minutes in the presence of 10% PEG-8000. Consistent with the microscopic phase regime, R175H also showed higher scattering than WT and R248Q in the presence of PEG, suggesting its higher propensity for LLPS (**Fig. 4e**). The liquid-like property of phase separated condensates were confirmed by the fusion (for WT condensates) (**Fig. 4f**) and high FRAP recovery (∼100%) with shorter t_1/2_ (∼<5s), immediately after their formation (0 h) (**Fig. 4g**, **Fig. S10a**). These condensates, however, showed reduced FRAP recovery with time, suggesting compromised dynamicity of the molecules inside the condensates, which is further promoted by both cancer-associated mutants (**Fig. 4g**). Further, the FTIR spectroscopy study of the dense and dilute fraction of WT p53C (separated by centrifugation) LLPS (at 0 h) showed no difference in their secondary structure after phase separation. (**Fig. 4h**).

**Figure 4.**
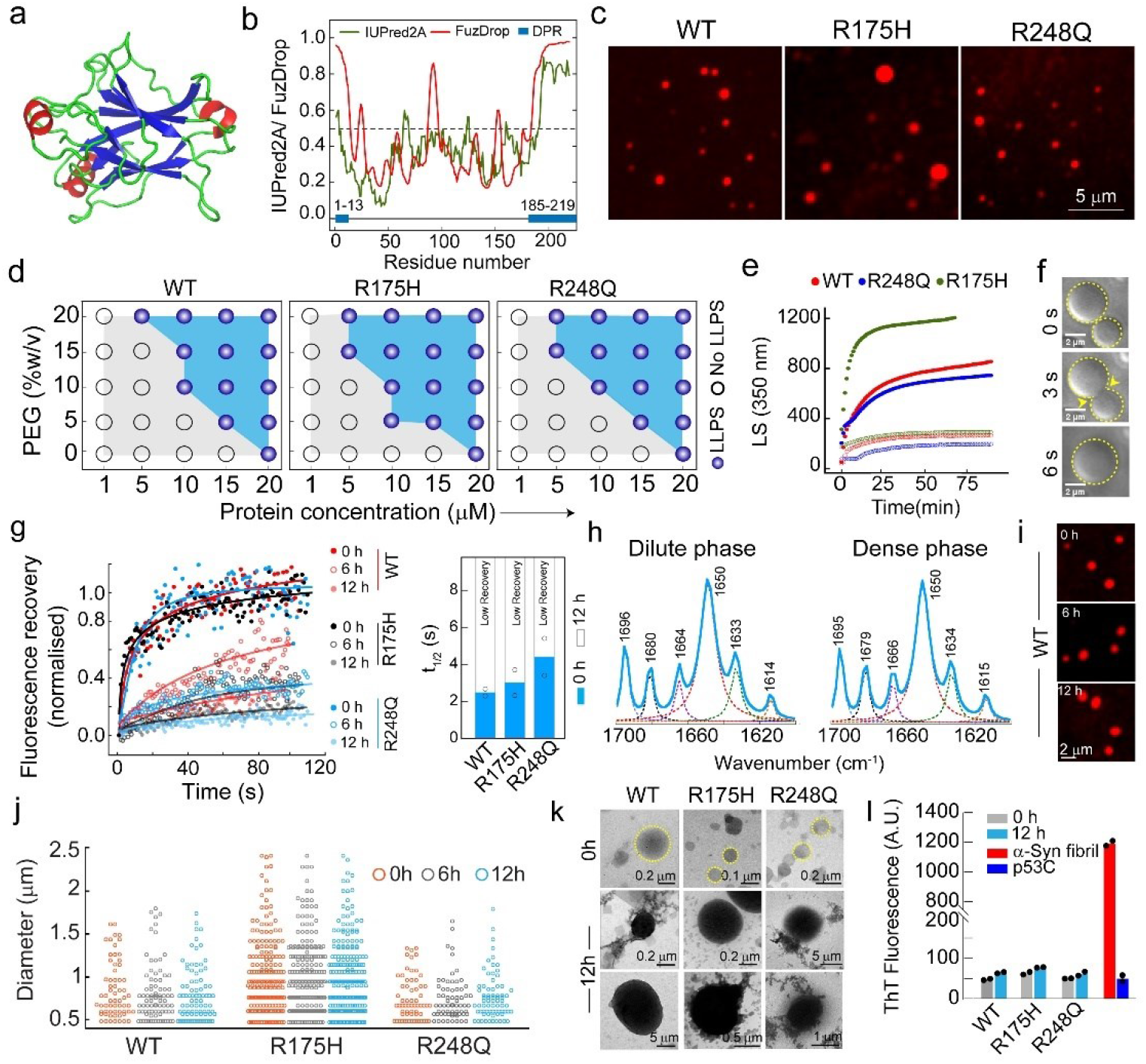
Liquid-liquid phase separation of p53C condensates *in vitro* and its transition from liquid to arrested state. **(a)** PDB structure of p53C (PDB ID: 2OCJ), showing the organization of its secondary structure (red represents α-helix, green represents random coil and blue represents β-sheet conformation). **(b)** *In silico* analysis of the primary sequence of WT p53 core protein (p53C) showing disorder tendency (green) using IUPRED2A, residue-based droplet promoting probabilities (red) and droplet-promoting regions (DPR) (blue box) using FuzDrop. **(c)** Microscopy images showing the condensate formation of NHS-Rhodamine labelled (10% v/v labelled to unlabelled protein) WT, R175H and R248Q p53C in the presence of 10% (w/v) polyethylene glycol 8000 (PEG-8000) in 50 mM sodium phosphate buffer (pH 7.4) (n=3 independent experiments). The scale bar is 5 μm. **(d)** Schematic representation of phase regime of WT, R175H, and R248Q showing the LLPS behaviour at different concentrations of protein and PEG-8000. Opaque circles represent LLPS, while open circles represent no LLPS. **(e)** Static light scattering (LS at 350 nm) showing condensate formation by WT p53C, R175H and R248Q in the presence (filled circles) and absence (open circles) of 10% PEG-8000. **(f)** Time-lapse DIC images showing the fusion event of two WT p53 condensates with time. The scale bar is 2 μm. **(g)** Fluorescence recovery after photobleaching (FRAP) experiment showing full FRAP recovery of freshly formed condensates but a reduction in recovery of all the three p53C variants with time. ***Right panel***: Estimation of t_1/2_ of the freshly formed condensates by WT and mutant p53C. Note: t_1/2_ for late time points was not estimated since no recovery was observed. **(h)** FTIR spectroscopy of dense and dilute phase of WT p53 after centrifugation showing no secondary structural change immediately after phase separation. **(i)** Representative confocal microscopy images of NHS-Rhodamine labelled [10% (v/v) labeled to unlabeled] 10 μM WT p53C condensates in the presence of 10% (w/v) PEG-8000 at different time intervals. Scale bar is 2 μm. **(j)** Size distribution of WT, R175H, and R248Q condensates with time showing the larger size of condensates for R175H compared to R248Q and WT. After formation, condensates by all these proteins, however, did not grow with time. **(k)** Representative TEM micrographs showing the morphology of condensates formed by 10 μM WT p53C, R175H and R248Q at 0 h. After 12 h, condensates show growth of aggregates from the condensates. **(l)** ThT binding assay showing no binding of ThT to aged (12 h) p53C condensates. α-Synuclein fibrils were used as a positive control. The experiments **(c, d and i)** were repeated three independents times and **(e-h, k and l)** were repeated two independent times.

Further, the time-dependent condensate formation/growth of WT p53C and its mutants suggests that the size and number of condensates did not change over time (up to 12 h) (**Fig. 4i, j and Fig. S10b**). However, with an increase in protein concentrations, although the condensate size did not alter, but the number of condensates increased, as shown for WT p53C (**Fig. S10c, d**). Further, the morphology of the condensates was characterized using transmission electron microscopy (TEM), which showed circular morphology at 0 h, which subsequently formed dark-round shape condensates and some amorphous aggregates appeared to originate from the condensates (**Fig. 4k**). However, the lack of ThT^88^ and CR^89^ dye (specific dye for amyloid aggregates) binding suggest that these aggregates might not be amyloid in nature (**Fig. 4l and Fig. S10e**). The data, therefore, suggest that p53C undergoes LLPS and liquid to arrested state transition with time, which is further promoted by both the cancer-associated p53 mutants.

### DNA/RNA modulates the formation and fate of multicomponent p53C LLPS

From the cellular studies, it was observed that the microenvironment of the nucleus and cytoplasm might dictate the fate and material properties of p53 condensates as nuclear p53 condensates maintain liquid-like property whereas cytoplasmic condensates rigidify with time **(Fig. 1 and 2)**. Since p53 is majorly known for its function as a transcription factor, which requires extensive interaction with nucleic acids in the cellular milieu, we studied the effect of nucleic acids on phase separation of p53. To examine this, p53C LLPS experiment was performed (with 10% PEG-8000) in the presence and absence of RNA, target DNA (TR-DNA, corresponding to the consensus sequence of the p53 response element for the p21 gene) and non-target DNA (NTR-DNA)^90^ (**Table S2**). Compared to condensates formed in the absence of any nucleic acid (p53C alone in the presence of 10% PEG-8000), the presence of both NTR-DNA and RNA increases the size and number of the p53 condensates (**Fig. 5a, b**). In contrast, in the presence of TR-DNA, p53C does not show a significant number of condensates, rather, a few smaller condensates were observed (**Fig. 5a, b)**. The dual fluorescence labelling study suggests that p53C condensates in the presence of NTR-DNA resulted in multicomponent phase separation (yellow condensate in the inset), while in the presence of TR-DNA, few residual DNA-protein conjugates were observed **(Fig. S11a, b)**, similar to previous study^91^. The facilitated p53C LLPS in the presence of RNA and NTR-DNA was further evident from the phase regime of p53C, which showed a decrease in the p53C concentration required for phase separation (**Fig. 5c, d and Fig. S11c, d**). Consistent with this, the LS experiment also showed enhanced LLPS kinetics of WT p53C in the presence of RNA and NTR-DNA but reduced LLPS kinetics in the presence of TR-DNA (**Fig. 5e**) compared to p53C alone sample. Interestingly, the NTR-DNA and RNA not only promote p53 phase separation but also help to maintain its liquid-like state as FRAP analysis showed >80% recovery at both 0 h and 12 h **(Fig. 5f and Fig. S11e),** in contrast to p53C alone sample where liquid-to-arrested state transition was observed **(Fig. 4g)**. Further, to directly probe the p53C condensates property in nuclear versus cytoplasmic microenvironment, we examined p53C condensate formation in nuclear (NE) and cytoplasmic extract (CE) of SaOS2 cells^92^ **(Fig. S12a, b)**. The data showed condensate formation of p53C in both extracts, however, with a lower concentration regime at 0 h **(Fig. S12c)**. Interestingly, the size of the p53C condensate in CE was higher, with lower circularity in comparison to condensate in NE **(Fig. 5g, h)**, indicating the rigidification of the condensates in CE. Indeed, the FRAP recovery of p53C condensates in NE was higher than the condensates formed in the CE **(Fig. 5i and Fig. S12d),** suggesting nuclear condensates remain in a relatively more liquid-like state, whereas the cytoplasmic microenvironment promotes rigidification of p53 condensates, consistent with our *in cellulo* data.

**Figure 5.**
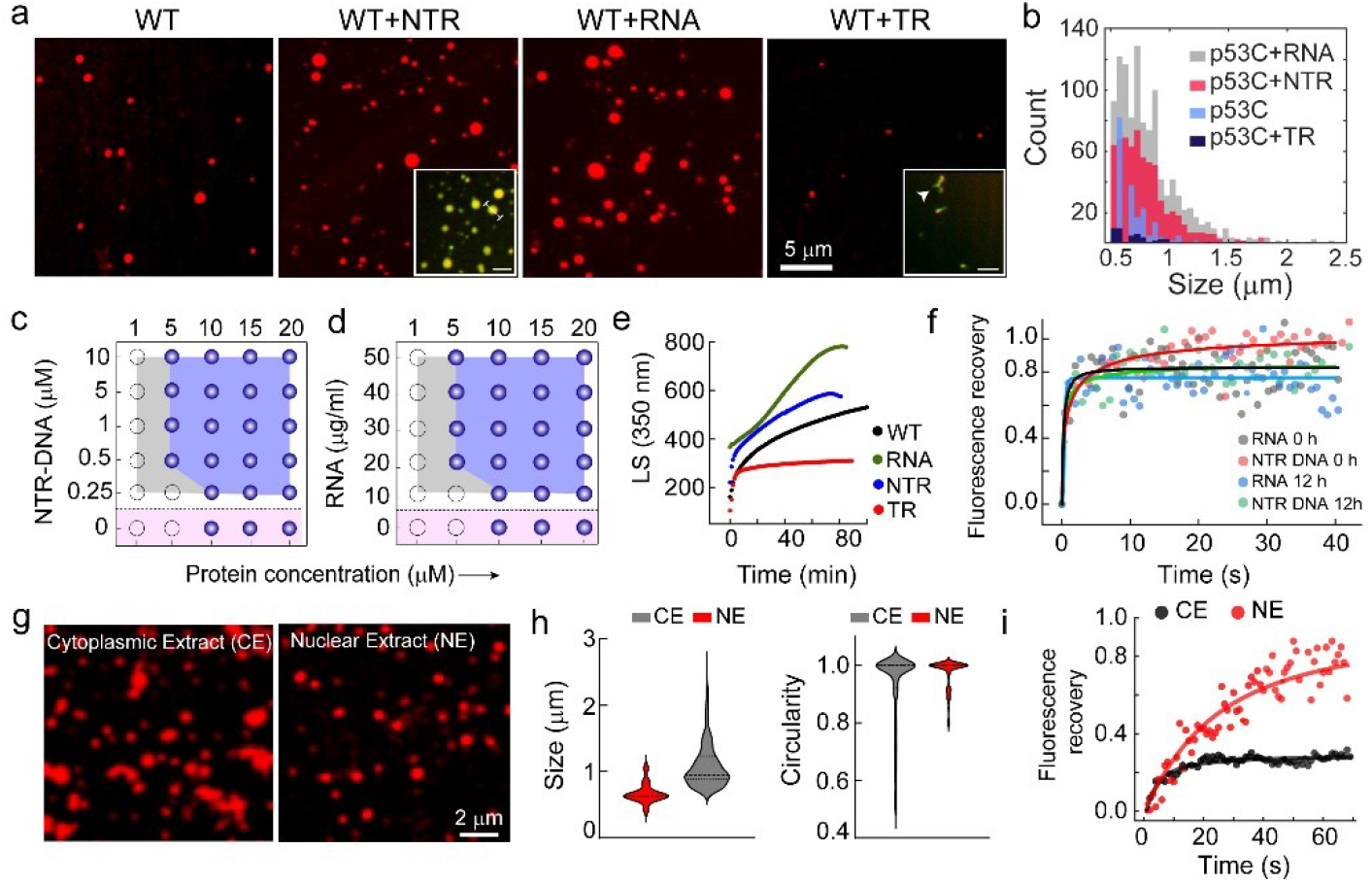
DNA and RNA modulate p53C phase separation. **(a)** Representative confocal images of WT p53C condensates formed in the presence and absence of target DNA (TR-DNA), non-target DNA (NTR-DNA) and RNA. The scale bar is 5 μm. The inset for NTR-DNA and TR-DNA represents the confocal microscopy images of 20 μM NHS-Rhodamine labelled [10% (v/v) labelled to unlabelled protein] p53C with Atto 488 labelled [10% (v/v) labelled to unlabelled DNA] NTR-DNA and TR-DNA showing multicomponent condensate formation. **(b)** Size (Feret) distributions of condensates formed by p53C alone and in the presence of TR-DNA, NTR-DNA and RNA. **(c-d)** Representation of phase regime of WT p53C [10% (v/v) NHS-Rhodamine labelled] in the presence of varying concentrations of NTR-DNA and RNA in 10% (w/v) PEG-8000 showing LLPS behaviour. Opaque circles represent LLPS and open circles represent no LLPS. Pink shaded region showing p53 condensates in the absence of any nucleic acid. **(e)** Static light scattering (LS at 350 nm) showing RNA and NTR-DNA promote p53C condensate formation, while the lesser extent of condensate in the presence of TR-DNA. **(f)** FRAP profile of p53C condensates in the presence of NTR-DNA and RNA showing a complete recovery at the early (0 h) and late (12 h) time points suggesting maintenance of the liquid-like state. **(g)** The confocal image of NHS-Rhodamine labelled p53C (10 μM) condensates formed in the cellular fraction of the nuclear extract (NE) and cytoplasmic extract (CE). The scale bar is 2 μm. **(h)** The size and circularity distribution of p53C condensates formed in the presence of CE and NE showing larger size in CE with lesser circularity in comparison to condensate formed in the NE. **(i)** FRAP recovery profile p53C condensates formed in NE showing a higher recovery profile (∼70%) compared to condensates formed in CE (∼20% FRAP recovery). The experiments **(a, c, d and g)** were repeated three independents times and **(e, f and i)** were repeated two independent times.

### Effect of target DNA (TR-DNA) on preformed p53C multicomponent condensates

Based on previous data, we hypothesize that the abundance of non-specific DNA and RNA in the nucleus might promote and stabilize the p53 liquid condensate, whereas cues due to exposure of specific DNA might promote their dissolution (Fig. 5). This de-mixing and mixing might be part of the p53 life cycle and function. In fact, when preformed WT p53C condensates were added on NTR-DNA coated coverslips, significantly more p53C condensates were seen. While with TR-DNA coated coverslips, we observed the major disappearance of the p53 preformed condensates **(Fig. 6a and Fig. S13a)**. This is further consistent with the p53 partitioning analysis in a single p53 condensate. The data suggest a significant reduction of p53 intensity in the dense phase when TR-DNA was added to preformed p53C condensate. In contrast, an increase in the dense phase signal was observed with the addition of NTR-DNA **(Fig. 6b and S13b, c)**. The de-mixing and mixing due to NTR-DNA versus TR-DNA were further confirmed by LS experiments, where after the formation of p53C condensates when RNA, TR-DNA, and NTR-DNA were added, the intensity values reduced significantly in the presence of TR-DNA, but slightly increased with the addition of NTR-DNA/ RNA (**Fig. 6c**).

**Figure 6.**
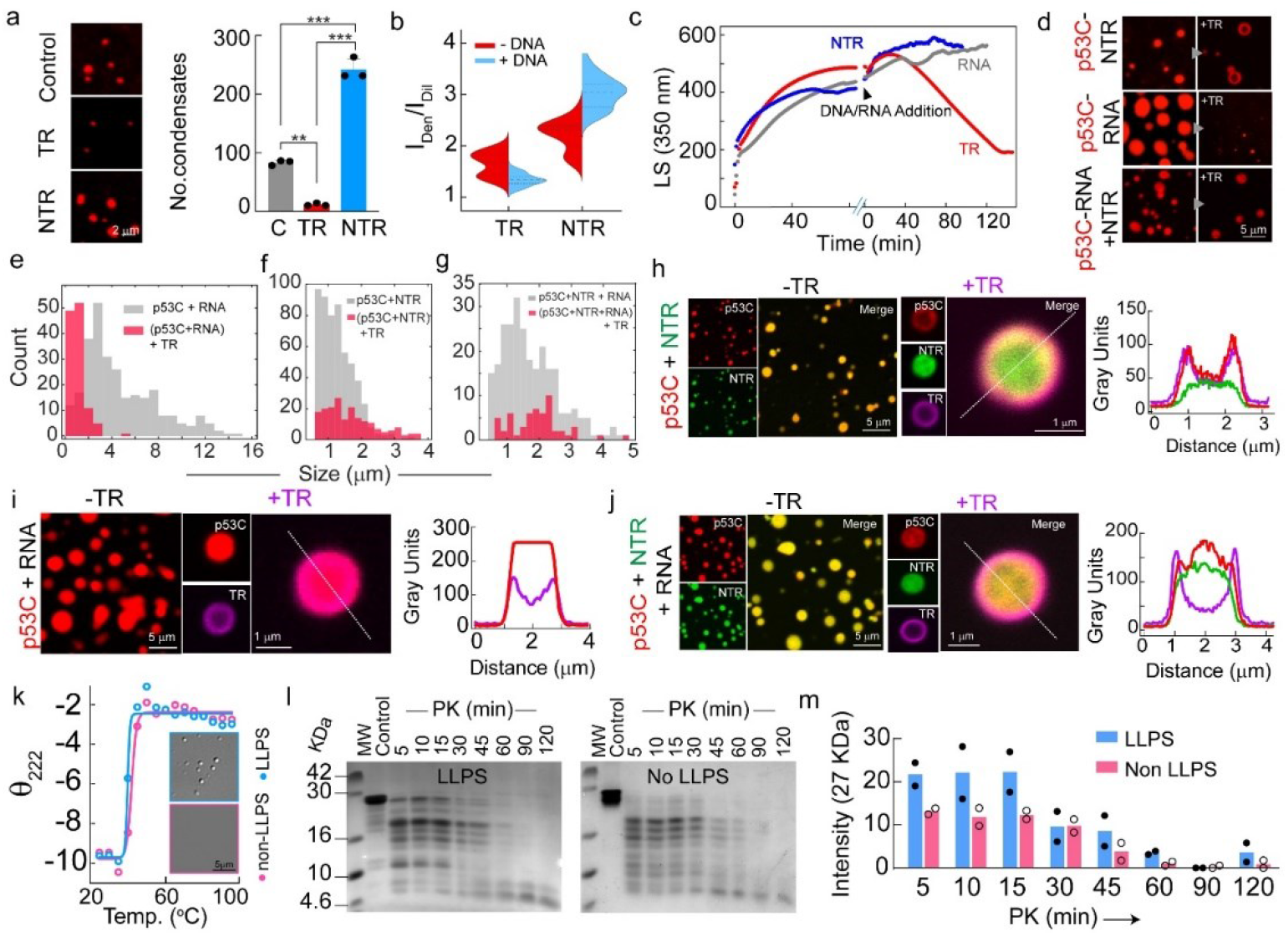
Effect of DNA/RNA on preformed p53C condensates. **(a) *Left panel:*** Representative images of preformed p53 condensates formed in the presence of 10% PEG-8000 after the addition onto the TR and NTR-DNA coated coverslips. Untreated coverslips were used as p53C control. (***Right panel***) The number of condensates with various nucleic acid coated surfaces showing an increased number of condensates in the presence of NTR-DNA while a significantly decreased number in the presence of TR-DNA as compared to the control (uncoated surface). p53C condensates formed in the presence of 10% PEG-8000 on uncoated coverslips were taken as control (C). The statistical significance was determined by one-way ANOVA using Newman−Keuls (SNK) post hoc analysis (p<0.001(***), p<0.01(**), p<0.05 (*)). **(b)** Split-violin plot representing the change in the partitioning of p53C after the addition of TR-DNA and NTR-DNA. **(c)** Static light scattering (LS at 350 nm) showing the kinetics of p53C in the presence of 10% (w/v) PEG-8000 and the effect post addition of TR-DNA, NTR-DNA and RNA. The data showing a decrease in light scattering value in the presence of TR-DNA, suggesting the dissolution of preformed p53C condensates. **(d)** The confocal microscopy images showing the effect of TR-DNA on preformed p53C condensate formed in the presence of NTR-DNA (p53C-NTR DNA), RNA (P53C-RNA) and RNA+NTR-DNA (p53-RNA-NTR-DNA) condensates. **(e-g)** Size distribution plot showing the reduction in the size/numbers of condensates after the addition of TR-DNA to **(e)** p53C-RNA condensates, **(f)** p53C-NTR-DNA condensates and **(g)** p53C-RNA-NTR-DNA condensates. **(h-j)** The confocal microscopy images showing few residual condensates that were observed after the dissolution of condensates due to the addition of TR-DNA. Left panel represents Rhod-p53 condensates formed in the presence of **(h)** Atto-488 labelled NTR DNA, **(i)** RNA and **(j)** RNA and Atto-488 labelled NTR-DNA. Middle panel represents residual p53C condensates that remained after the addition of Atto-647N labelled TR-DNA. Right panel represents a line profile across the condensates to show the spatial distribution of protein and DNA. **(k)** The change in molar ellipticity at θ_222_ with an increase in temperature showing no major changes in temperature denaturing profiles between the LLPS and non-LLPS state of p53C. Representative microscopic images of p53C in LLPS and non-LLPS states during the temperature-dependent CD experiment are shown. The scale bar is 5 μm. **(l)** Proteinase K (PK) digestion followed by SDS-PAGE of WT p53C LLPS and non-LLPS state showing higher PK resistivity of p53 LLPS state. **(m)** SDS-PAGE quantification of PK digestion assay of LLPS and non-LLPS p53C showing faster PK digestion with time of p53C in non-LLPS state compared to LLPS state. The experiments **(a, c, d and h-l)** were repeated two independent times.

We further examined the effect of TR-DNA on multicomponent p53 condensates formed in the presence of NTR-DNA (p53C-NTR-DNA) and RNA (p53C-RNA). The addition of TR-DNA to preformed p53C-RNA condensates led to the dissolution of p53C-RNA condensates. However, few condensates with smaller size were seen **(Fig. 6d, e)**. Similarly, the addition of TR-DNA to preformed p53C-NTR-DNA and p53C-NTR-DNA+RNA multicomponent condensates resulted in the dissolution of most of the preformed p53 condensates as evident from the reduced number and size of the condensates **(Fig. 6d, f and g)**. Interestingly, the residual p53C-NTR-DNA condensates present after the addition of TR-DNA resulted in two distinct morphologies, one is spherical and the other one is hollow-ring-like condensates **(Fig. 6d)**.

To further characterise the spatial distribution of protein and nucleic acids in the multicomponent system, we performed the fluorescence study using Atto 647N-labelled TR-DNA, Atto 488-labelled NTR-DNA and NHS-Rhodamine-labelled p53C. For residual NTR-p53C condensates (which are large enough to resolve spatially), we observed that TR-DNA and p53C both partitioned at the periphery of the condensate, while NTR-DNA is localized majorly at the center of the condensate **(Fig. 6h and Fig. S13d)**. Similar results were also obtained when TR-DNA was added to p53C-RNA and NTR-RNA-p53C multicomponent condensates, resulting in a few residual condensates, where TR-DNA remained at the periphery and p53C localized at the center of the condensates **(Fig. 6i, j and Fig S13d, e)**.

We hypothesize that the binding strength of nucleic acids with p53 might dictate the mixed versus de-mixed state. Indeed, our binding study using spectral-shift assay^93^ suggests strong binding of p53 with specific DNA over non-specific DNA **(Fig. S13f, g)**. Therefore, stronger bindings to specific DNA dissolved the preformed condensate, which could be due to the specific interaction between TR-DNA and p53C in addition to general protein-nucleic acid electrostatic interactions.

To address whether p53 condensate formation has any evolutionary advantages over soluble state, we examined the thermal stability assay using DSC and CD spectroscopy. Both melting temperature by DSC and ϴ_222_ by CD showed no difference between p53C LLPS and non-LLPS state **(Fig. 6k and Fig. S14)**. However, p53C LLPS showed higher proteinase K (PK) resistivity, as shown in **Fig. 6 (l-m)**, compared to non-LLPS p53. This data suggests that p53 condensate possesses an advantage not only as a ready pool for immediate action (p53 DNA binding for apoptosis, cell cycle arrest when needed) but also provides protease resistivity, countering their cellular instability.

## Discussion

LLPS of biomolecules (such as proteins and nucleic acids) has been implicated in driving the formation of membraneless compartments in cells, commonly designated as biomolecular condensates^5,6,10–12,94^. These liquid-like condensates are quickly responsive to immediate environmental changes and thus perform various cellular functions^17,95–100^. Recent data suggested that the cell nucleus is not homogeneous in nature but rather consists of a heterogeneous mixture of subnuclear membraneless compartments capable of performing different biological processes^33,101,102^. These membraneless organizations include Cajal body, nuclear speckles, nucleoli, etc^5,6,16,57^. In this context, many transcription factors are known to be heterogeneously localized in the nucleus, which are indeed in biomolecular condensate state^15,103–106^. Further, the nuclear condensates consisting of transcription factors and coactivators are shown to regulate the transcription process of key genes associated with cellular identity and functionality^15,33,106–109^.

p53 is a tumor suppressor protein and transcription factor, which generally maintains a low-level concentration in normal cells^69,110^. However, upon stress signals such as DNA damage, p53 gets phosphorylated and translocated to the nucleus, where it is concentrated and binds to DNA. This event is responsible for a plethora of activities regulating tumor suppressor functions such as DNA repair, cell cycle arrest, or apoptosis^70,111^. Mutation of p53 not only results in the loss of tumor suppressive function but also results in their gain-of-oncogenic function by inactivating WT p53 and other tumor suppressors, known as “dominant negative effect” ^45^. Recently, it was suggested that p53 aggregation and amyloid formation may also be linked with tumorigenic potential in cells^48,112^. Previous study has also indicated that p53 aggregation might be associated with p53 LLPS, where liquid-to-solid-like transition leads to amyloid fibril formation^53–55^, similar to many other neurodegenerative disorders^5,29,31,113^. Further, p53 condensate formation is also shown to be modulated by post-translational modifications^114^. However, the mechanism and role of various nuclear constituents (such as RNA and DNA) on spatio-temporal p53 phase separation in cells are not known. Further, it is not yet clear whether p53 condensates are functional and how their material properties might be linked with cancer pathogenesis. In this context, several studies have shown the formation of biomolecular condensates by cancer-associated proteins such as SPOP^115^, BRD4^116^, and fusion oncoproteins^117,118^.

In our current work, we have investigated the condensate formation ability of p53 in mammalian cells, followed by a molecular-level investigation using *in vitro* approach. We showed that once p53 gets stabilized in the cancer cells or is overexpressed in cells, p53 is heterogeneously concentrated in the nucleus **(Fig. 1d)**, as shown for other transcription factors^4,15,33,106^. After overexpression in cells, initially, p53 was shown to be localized inside the nucleus as a liquid-like condensate, but with time, it partially translocated to the cytoplasm and underwent LLPS, which transitioned into an arrested state. In contrast, the nuclear p53 condensates remained liquid-like (high FRAP recovery and condensate fusion) with time **(Fig. 1k and 2j)**. The nuclear to cytoplasmic partitioning and subsequent solidification are also shown for other transcription factors such as ARF^119^. This data indicates that liquid-like p53 condensates in the nucleus might be functional (transcriptionally active), whereas cytoplasmic condensates, which become arrested over time, could be degraded for overall cellular fitness. Indeed, p53 mutations not only led to a more arrested state of the condensates in the nucleus but also fastened the liquid-to-arrested state transition in the cytoplasm, suggesting a non-functional state of mutants. This is further supported by staining of misfolded p53, where mutant p53 condensates showed a higher degree of misfolding both in the nucleus and cytoplasm in contrast to WT protein **(Fig. 2k and l)**. Therefore, mutations promote p53 misfolding in the condensate state, and their arrested state transition could be associated with p53 loss of function, which is indeed shown by the loss of DNA binding **(Fig. 3d, e)**. In this context, it was shown previously that transcription factors also undergo a liquid-to-arrested state, where this transition results in the loss of transcriptional activity^120^.

It has been shown that transcription factors bind to various nuclear components, including nuclear RNA and/or DNA, which promote and modulate the condensate state and their transcriptional activity^4,25,33,91,121–124^. Further, nuclear RNAs are also known to participate in multicomponent phase separation with various proteins for membraneless organelle formation in the nucleus, such as nucleoli and nuclear bodies, including Cajal bodies^16,33,125,126^. To understand the role of different nucleic acids in the formation of p53 condensate and its material properties, p53C LLPS was studied *in vitro*. Our data showed that p53C readily undergoes LLPS upon incubation **(Fig. 4c)**, and with time, it transforms into an arrested state **(Fig. 4g)**. The LLPS and subsequent rigidification of p53C are further fastened by cancers-associated hotspot mutations. Interestingly, both RNA and non-specific DNA not only enhanced the p53C phase separation but also helped to maintain its liquid-like nature **(Fig. 5a, f)**. RNA and non-specific DNA might template or promote the multivalent protein-protein, protein DNA/RNA interaction for p53C phase separation^23^. These data are consistent with our *in cellulo* data of p53 condensates. Interestingly, the presence of specific DNA not only inhibits the p53 de-mixing but also dissolves the preformed p53 condensates (formed either by p53C alone or in the presence of DNA and/or RNA) **(Fig. 6)**. Previous study also suggests that nucleic acids modulate the formation of multicomponent LLPS and material property of the condensates^23,104,127^. Our data thus suggests that upon entry into the nucleus, p53 readily undergoes LLPS into a micro-phase separated condensate state^79,128^ in the presence of nucleic acid, which might act as a ready pool of functional p53. This process might help p53 to stay in the vicinity of specific DNA (upon exposure)^129^, which subsequently may get dissolved into a soluble state for performing the tumor suppressor/transcriptional activity. This dissolution upon contact with specific DNA could also be linked with the wetting phenomenon of p53 condensates on DNA^91^ **(Fig. S15)**.

Further, the exposure of p53 target genes might also direct the movement of p53 condensates, where p53 becomes a mixed state for attaining higher thermodynamic stability. In this direction, it was shown that droplets containing urease move towards the gradient of urea, resulting in a mixed state^130^. Further, once the p53 load increases in the nucleus, some nuclear p53 population is transported into the cytoplasm through the nuclear pore complex, where it undergoes LLPS and subsequently solidifies with time. However, at this stage, whether this solidification by WT or mutant is associated with p53 amyloid formation is not clear, as OC staining did not indicate the presence of p53 amyloids.

Further, the p53 phase separation not only enhances the protein fitness against its degradation but also the local concentration of p53 may increase the availability of the protein to function as a transcription factor. The LLPS of p53 and its mis-regulated transition to an arrested state followed by misfolding might result in the loss of tumor suppressive and gain of oncogenic function, which may be linked with cancer.

## Methods

The *in cellulo* phase separation studies of p53 and its hotspot mutants (R175H and R248Q) were performed in GFP-p53 transfected HeLa and SaOS2 cell lines. The nuclear and cytoplasmic condensate formation was examined using confocal, STED and lattice light-sheet microscopy. The dynamic nature of the condensates was characterized through FRAP and fusion experiments. The p53 expression in the cells was monitored over time using Western blot analysis. The *in vitro* LLPS of WT p53 core (p53C) and mutants were performed in the presence of 50 mM sodium phosphate (buffer pH 7.4) and different concentrations of PEG-8000. The condensates were observed under the confocal microscope. The change in secondary structural conformation of p53 due to LLPS was examined using FTIR spectroscopy. The aggregate formation during LLPS was studied using ThT fluorescence and Congo Red binding assay, and the morphology was analyzed using TEM. The transcriptional activity and DNA binding ability of p53 against p21-specific DNA were determined by ELISA and ChIP experiments, respectively. The gene expression of p21 was monitored by qRT-PCR analysis. Apoptotic cell death of p53 WT and mutants at different time points was studied by Annexin-PI assay followed by FACS analysis. The detailed methods are provided in Supplementary Information.

## Data Availability

All the data in this manuscript is available in the main and supplementary information. The tools used for plotting and analysis have been mentioned in the main and supplementary manuscript.

## Acknowledgments

The authors thank the IIT Bombay central facilities, namely, microscopy and FACS facility at BSBE, TEM and FTIR facilities at SAIF. The authors also thank Prof. Ranjith Padinhateeri, BSBE, IIT Bombay, for valuable suggestions. The authors also thank Dr. Nitu Singh, RCB Faridabad, Santosh Panigrahi, Pradip Shinde and Dr. Vidhu Soman from microscopy facility at IIT Bombay, Vijay Vittal, Technical Assistant for microscopy facility at IISER Pune, Dr. Saji Menon, Senior Field Application Scientist, and Dr. Ruchika Dadhich, Application Scientist at Nanotemper Technologies, Dr. Debdeep Chatterjee for valuable inputs in experiments. The authors also thank DST-SERB (File no. CRG/2019/001133) and TATA Innovation (BT/HRD/35/01/03/2020) for financial support. S.S. acknowledges the DBT/Wellcome Trust India Alliance Fellowship [IA/E/17/1/503663] for financial support. D.D. acknowledges MoE, Govt. of India, for the GATE fellowship.

## Author Contribution

The study was conceptualized by S.K.M. and designed by S.K.M., D.D. Under the guidance of K.S., D.D. and A.N. performed STED experiments. D.D. performed *in silico* analysis. D.D., A.N., A.S., A.P., K.P., S.M, S.S., S.P., N.G. performed *in cellulo* experiments. D.D., A.N., A.P., K.P., L.G., M.P., J.D., A.S.S., R.S., and S. Mukherjee performed *in vitro* experiments. P.K. M.P. and D.D. prepared the illustration. S.K.M., D.D. and A.N. participated in writing the manuscript. S.K.M., D.D., A.N., A.S., K.P. and S. Mukherjee edited the draft. All the authors discussed and approved the final version of the manuscript.

## References

1 Friedman, J. R. & Nunnari, J. Mitochondrial form and function. Nature 505, 335–343 (2014).

2 Luzio, J. P., Pryor, P. R. & Bright, N. A. Lysosomes: fusion and function. Nature reviews Molecular cell biology 8, 622–632 (2007).

3 Boisvert, F.-M., van Koningsbruggen, S., Navascués, J. & Lamond, A. I. The multifunctional nucleolus. Nature reviews Molecular cell biology 8, 574–585 (2007).

4 Banani, S. F. et al. Compositional Control of Phase-Separated Cellular Bodies. Cell 166, 651–663 (2016).

5 Shin, Y. & Brangwynne, C. P. Liquid phase condensation in cell physiology and disease. Science 357, eaaf4382 (2017).

6 Banani, S. F., Lee, H. O., Hyman, A. A. & Rosen, M. K. Biomolecular condensates: organizers of cellular biochemistry. Nature reviews Molecular cell biology 18, 285–298 (2017).

7 Woodruff, J. B., Hyman, A. A. & Boke, E. Organization and function of non-dynamic biomolecular condensates. Trends in biochemical sciences 43, 81–94 (2018).

8 Hyman, A. A. & Simons, K. Beyond oil and water—phase transitions in cells. Science 337, 1047–1049 (2012).

9 Hyman, A. A., Weber, C. A. & Julicher, F. Liquid-liquid phase separation in biology. Annu Rev Cell Dev Biol 30, 39–58 (2014).

10 Lyon, A. S., Peeples, W. B. & Rosen, M. K. A framework for understanding the functions of biomolecular condensates across scales. Nature Reviews Molecular Cell Biology 22, 215–235 (2021).

11 Alberti, S. Phase separation in biology. Current Biology 27, R1097–R1102 (2017).

12 Boeynaems, S. et al. Protein phase separation: a new phase in cell biology. Trends in cell biology 28, 420–435 (2018).

13 Alberti, S., Gladfelter, A. & Mittag, T. Considerations and Challenges in Studying Liquid-Liquid Phase Separation and Biomolecular Condensates. Cell 176, 419–434 (2019).

14 Lu, H. et al. Phase-separation mechanism for C-terminal hyperphosphorylation of RNA polymerase II. Nature 558, 318–323 (2018).

15 Hnisz, D., Shrinivas, K., Young, R. A., Chakraborty, A. K. & Sharp, P. A. A phase separation model for transcriptional control. Cell 169, 13–23 (2017).

16 Fox, A. H., Nakagawa, S., Hirose, T. & Bond, C. S. Paraspeckles: where long noncoding RNA meets phase separation. Trends in biochemical sciences 43, 124–135 (2018).

17 Riback, J. A. et al. Stress-triggered phase separation is an adaptive, evolutionarily tuned response. Cell 168, 1028–1040 (2017).

18 Cramer, P. Organization and regulation of gene transcription. Nature 573, 45–54 (2019).

19 Kribelbauer, J. F., Rastogi, C., Bussemaker, H. J. & Mann, R. S. Low-affinity binding sites and the transcription factor specificity paradox in eukaryotes. Annual review of cell and developmental biology 35, 357–379 (2019).

20 Calabretta, S. & Richard, S. Emerging roles of disordered sequences in RNA-binding proteins. Trends in Biochemical Sciences 40, 662–672 (2015).

21 Martin, E. W. et al. Valence and patterning of aromatic residues determine the phase behavior of prion-like domains. Science 367, 694–699 (2020).

22 Pak, C. W. et al. Sequence Determinants of Intracellular Phase Separation by Complex Coacervation of a Disordered Protein. Molecular Cell 63, 72–85 (2016).

23 Maharana, S. et al. RNA buffers the phase separation behavior of prion-like RNA binding proteins. Science 360, 918–921 (2018).

24 Bremer, A. et al. Deciphering how naturally occurring sequence features impact the phase behaviours of disordered prion-like domains. Nature Chemistry 14, 196–207 (2022).

25 Lin, Y., Protter, D. S. W., Rosen, M. K. & Parker, R. Formation and Maturation of Phase-Separated Liquid Droplets by RNA-Binding Proteins. Molecular Cell 60, 208–219 (2015).

26 Wang, J. et al. A molecular grammar governing the driving forces for phase separation of prion-like RNA binding proteins. Cell 174, 688–699 (2018).

27 Saha, S. et al. Polar positioning of phase-separated liquid compartments in cells regulated by an mRNA competition mechanism. Cell 166, 1572–1584 (2016).

28 Brangwynne, C. P., Tompa, P. & Pappu, R. V. Polymer physics of intracellular phase transitions. Nature Physics 11, 899–904 (2015).

29 Alberti, S. & Hyman, A. A. Biomolecular condensates at the nexus of cellular stress, protein aggregation disease and ageing. Nature Reviews Molecular Cell Biology 22, 196–213 (2021).

30 Jiang, S., Fagman, J. B., Chen, C., Alberti, S. & Liu, B. Protein phase separation and its role in tumorigenesis. eLife 9, e60264 (2020).

31 Alberti, S. & Dormann, D. Liquid-Liquid Phase Separation in Disease. Annual Review of Genetics 53, 171–194 (2019).

32 Wei, M.-T. et al. Nucleated transcriptional condensates amplify gene expression. Nature Cell Biology 22, 1187–1196 (2020).

33 Bhat, P., Honson, D. & Guttman, M. Nuclear compartmentalization as a mechanism of quantitative control of gene expression. Nature Reviews Molecular Cell Biology 22, 653–670 (2021).

34 Levine, A. J. p53, the cellular gatekeeper for growth and division. Cell 88, 323–331 (1997).

35 Fuster, J. J., Sanz-González, S. M., Moll, U. M. & Andrés, V. Classic and novel roles of p53: prospects for anticancer therapy. Trends in molecular medicine 13, 192–199 (2007).

36 Zilfou, J. T. & Lowe, S. W. Tumor suppressive functions of p53. Cold Spring Harbor perspectives in biology 1, a001883 (2009).

37 Levine, A. J., Momand, J. & Finlay, C. A. The p53 tumour suppressor gene. Nature 351, 453–456 (1991).

38 Harms, K., Nozell, S. & Chen, X. The common and distinct target genes of the p53 family transcription factors. Cellular and Molecular Life Sciences 61, 822–842 (2004).

39 Vousden, K. H. & Lu, X. Live or let die: the cell’s response to p53. Nature Reviews Cancer 2, 594–604 (2002).

40 Kruiswijk, F., Labuschagne, C. F. & Vousden, K. H. p53 in survival, death and metabolic health: a lifeguard with a licence to kill. Nature Reviews Molecular Cell Biology 16, 393–405 (2015).

41 Bieging, K. T., Mello, S. S. & Attardi, L. D. Unravelling mechanisms of p53-mediated tumour suppression. Nature Reviews Cancer 14, 359–370 (2014).

42 Moll, U. M., LaQuaglia, M., Bénard, J. & Riou, G. Wild-type p53 protein undergoes cytoplasmic sequestration in undifferentiated neuroblastomas but not in differentiated tumors. Proceedings of the National Academy of Sciences of the U.S.A. 92, 4407–4411 (1995).

43 Moll, U. M. et al. Cytoplasmic sequestration of wild-type p53 protein impairs the G1 checkpoint after DNA damage. Molecular and cellular biology 16, 1126–1137 (1996).

44 Moll, U. M., Riou, G. & Levine, A. J. Two distinct mechanisms alter p53 in breast cancer: mutation and nuclear exclusion. Proceedings of the National Academy of Sciences of the U.S.A. 89, 7262–7266 (1992).

45 Xu, J. et al. Gain of function of mutant p53 by coaggregation with multiple tumor suppressors. Nature chemical biology 7, 285–295 (2011).

46 De Smet, F. et al. Nuclear inclusion bodies of mutant and wild-type p53 in cancer: a hallmark of p53 inactivation and proteostasis remodelling by p53 aggregation. The Journal of pathology 242, 24–38 (2017).

47 Ghosh, S. et al. p53 amyloid formation leading to its loss of function: implications in cancer pathogenesis. Cell Death & Differentiation 24, 1784–1798 (2017).

48 Navalkar, A. et al. Direct evidence of cellular transformation by prion-like p53 amyloid infection. Journal of Cell Science 134, jcs258316 (2021).

49 Navalkar, A. et al. Prion-like p53 amyloids in cancer. Biochemistry 59, 146–155 (2019).

50 Joerger, A. C. & Fersht, A. R. The p53 pathway: origins, inactivation in cancer, and emerging therapeutic approaches. Annual review of biochemistry 85, 375–404 (2016).

51 Silva, J. L., Gallo, C. V. D. M., Costa, D. C. & Rangel, L. P. Prion-like aggregation of mutant p53 in cancer. Trends in biochemical sciences 39, 260–267 (2014).

52 Costa, D. C. et al. Aggregation and prion-like properties of misfolded tumor suppressors: is cancer a prion disease? Cold Spring Harbor perspectives in biology 8, a023614 (2016).

53 Safari, M. S. et al. Anomalous dense liquid condensates host the nucleation of tumor suppressor p53 fibrils. Iscience 12, 342–355 (2019).

54 Yang, D. S. et al. Mesoscopic protein-rich clusters host the nucleation of mutant p53 amyloid fibrils. Proceedings of the National Academy of Sciences of the U.S.A. 118, e2015618118 (2021).

55 Petronilho, E. C. et al. Phase separation of p53 precedes aggregation and is affected by oncogenic mutations and ligands. Chemical Science 12, 7334–7349 (2021).

56 Guo, A. et al. The function of PML in p53-dependent apoptosis. Nature Cell Biology 2, 730–736 (2000).

57 Cioce, M. & Lamond, A. I. in Annual Review of Cell and Developmental Biology 21, 105–131 (2005).

58 Cannon, J. V. & Lane, D. P. Protein synthesis required to anchor a mutant p53 protein which is temperature-sensitive for nuclear transport. Nature 349, 802–806 (1991).

59 Shaulsky, G., Goldfinger, N., Peled, A., & Rotter, V. Involvement of wild-type p53 protein in the cell cycle requires nuclear localization. Cell growth & differentiation: the molecular biology journal of the American Association for Cancer Research 2, 661–667 (1991).

60 Oren, M., Maltzman, W. & Levine, A. J. Post-translational regulation of the 54K cellular tumor antigen in normal and transformed cells. Molecular and cellular biology 1, 101–110 (1981).

61 Slade, N. & Moll, U. M. Mutational analysis of p53 in human tumors: immunocytochemistry. p53 Protocols 234, 231–243 (2003).

62 Meek, D. W. The p53 response to DNA damage. DNA repair 3, 1049–1056 (2004).

63 Joerger, A. C. & Fersht, A. R. Structural biology of the tumor suppressor p53. Annual Review of Biochemistry. 77, 557–582 (2008).

64 Miskei, M., Horvath, A., Vendruscolo, M. & Fuxreiter, M. Sequence-based prediction of fuzzy protein interactions. Journal of molecular biology 432, 2289–2303 (2020).

65 Jumper, J. et al. Highly accurate protein structure prediction with AlphaFold. Nature 596, 583–589 (2021).

66 Solares, M. J. et al. High-Resolution Imaging of Human Cancer Proteins Using Microprocessor Materials. ChemBioChem 23, e202200310 (2022).

67 Maki, C. G., Huibregtse, J. M. & Howley, P. M. In vivo ubiquitination and proteasome-mediated degradation of p53. Cancer research 56, 2649–2654 (1996).

68 Moll, U. M. & Petrenko, O. The MDM2-p53 interaction. Molecular cancer research 1, 1001–1008 (2003).

69 Kubbutat, M. H., Jones, S. N. & Vousden, K. H. Regulation of p53 stability by Mdm2. Nature 387, 299–303 (1997).

70 Vogelstein, B., Lane, D. & Levine, A. J. Surfing the p53 network. Nature 408, 307–310 (2000).

71 Hu, J., Lieb, J. D., Sancar, A. & Adar, S. Cisplatin DNA damage and repair maps of the human genome at single-nucleotide resolution. Proceedings of the National Academy of Sciences of the U.S.A. 113, 11507–11512 (2016).

72 Blom, H. & Widengren, J. Stimulated emission depletion microscopy. Chemical reviews 117, 7377–7427 (2017).

73 Hell, S. W. & Wichmann, J. Breaking the diffraction resolution limit by stimulated emission: stimulated-emission-depletion fluorescence microscopy. Optics Letters 19, 780–782 (1994).

74 Ottaviano, L. et al. Molecular characterization of commonly used cell lines for bone tumor research: a trans-European EuroBoNet effort. Genes, Chromosomes and Cancer 49, 40–51 (2010).

75 Chen, X., Ko, L. J., Jayaraman, L. & Prives, C. p53 levels, functional domains, and DNA damage determine the extent of the apoptotic response of tumor cells. Genes & development 10, 2438–2451 (1996).

76 Kudo, N. et al. Leptomycin B inactivates CRM1/exportin 1 by covalent modification at a cysteine residue in the central conserved region. Proceedings of the National Academy of Sciences of the U.S.A. 96, 9112–9117 (1999).

77 Greenspan, P., Mayer, E. P. & Fowler, S. D. Nile red: a selective fluorescent stain for intracellular lipid droplets. The Journal of cell biology 100, 965–973 (1985).

78 Choi, J.-M., Holehouse, A. S. & Pappu, R. V. Physical principles underlying the complex biology of intracellular phase transitions. Annual review of biophysics 49, 107–133 (2020).

79 Kar, M. et al. Phase-separating RNA-binding proteins form heterogeneous distributions of clusters in subsaturated solutions. Proceedings of the National Academy of Sciences of the U.S.A. 119, e2202222119 (2022).

80 Choi, J.-M., Hyman, A. A. & Pappu, R. V. Generalized models for bond percolation transitions of associative polymers. Physical Review E 102, 042403 (2020).

81 Stephen, C. W. & Lane, D. P. Mutant conformation of p53: precise epitope mapping using a filamentous phage epitope library. Journal of molecular biology 225, 577–583 (1992).

82 Brandt, T., Kaar, J. L., Fersht, A. R. & Veprintsev, D. B. Stability of p53 homologs. PLoS One 7, e47889 (2012).

83 Bullock, A. N. et al. Thermodynamic stability of wild-type and mutant p53 core domain. Proceedings of the National Academy of Sciences of the U.S.A. 94, 14338–14342 (1997).

84 Ang, H. C., Joerger, A. C., Mayer, S. & Fersht, A. R. Effects of common cancer mutations on stability and DNA binding of full-length p53 compared with isolated core domains. Journal of Biological Chemistry 281, 21934–21941 (2006).

85 Wang, Y., Rosengarth, A. & Luecke, H. Structure of the human p53 core domain in the absence of DNA. Acta Crystallographica Section D: Biological Crystallography 63, 276–281 (2007).

86 Mészáros, B., Erdős, G. & Dosztányi, Z. IUPred2A: context-dependent prediction of protein disorder as a function of redox state and protein binding. Nucleic Acids Research 46, W329–W337 (2018).

87 Erdős, G. & Dosztányi, Z. Analyzing protein disorder with IUPred2A. Current Protocols in Bioinformatics 70, e99 (2020).

88 LeVine III, H. Quantification of β-sheet amyloid fibril structures with thioflavin T. Methods in enzymology 309, 274–284 (1999).

89 Klunk, W. E., Jacob, R. F. & Mason, R. P. Quantifying amyloid by congo red spectral shift assay. Methods in enzymology 309, 285–305 (1999).

90 Kamagata, K. et al. Liquid-like droplet formation by tumor suppressor p53 induced by multivalent electrostatic interactions between two disordered domains. Scientific reports 10, 1–12 (2020).

91 Morin, J. A. et al. Sequence-dependent surface condensation of a pioneer transcription factor on DNA. Nature Physics 18, 271–276 (2022).

92 Poudyal, M. et al. Intermolecular interactions underlie protein/peptide phase separation irrespective of sequence and structure at crowded milieu. Nature Communications 14, 6199 (2023).

93 Langer, A. et al. A New Spectral Shift-Based Method to Characterize Molecular Interactions. ASSAY and Drug Development Technologies 20, 83–94 (2022).

94 Hyman, A. A., Weber, C. A. & Jülicher, F. in Annual review of cell and developmental biology 30, 39–58 (2014).

95 Franzmann, T. M. et al. Phase separation of a yeast prion protein promotes cellular fitness. Science 359, eaao5654 (2018).

96 Hubstenberger, A. et al. P-body purification reveals the condensation of repressed mRNA regulons. Molecular cell 68, 144–157 (2017).

97 Sheu-Gruttadauria, J. & MacRae, I. J. Phase transitions in the assembly and function of human miRISC. Cell 173, 946–957 (2018).

98 Brangwynne, C. P. et al. Germline P granules are liquid droplets that localize by controlled dissolution/condensation. Science 324, 1729–1732 (2009).

99 Mitrea, D. M. & Kriwacki, R. W. Phase separation in biology; functional organization of a higher order. Cell Communication and Signaling 14, 1–20 (2016).

100 Franzmann, T. M. & Alberti, S. Protein phase separation as a stress survival strategy. Cold Spring Harbor perspectives in biology 11, a034058 (2019).

101 Strom, A. R. et al. Phase separation drives heterochromatin domain formation. Nature 547, 241–245 (2017).

102 Sabari, B. R., Dall’Agnese, A. & Young, R. A. Biomolecular condensates in the nucleus. Trends in biochemical sciences 45, 961–977 (2020).

103 Boija, A. et al. Transcription factors activate genes through the phase-separation capacity of their activation domains. Cell 175, 1842–1855 (2018).

104 Gibson, B. A. et al. Organization of chromatin by intrinsic and regulated phase separation. Cell 179, 470–484 (2019).

105 Lu, Y. et al. Phase separation of TAZ compartmentalizes the transcription machinery to promote gene expression. Nature cell biology 22, 453–464 (2020).

106 Kimura, H. & Sato, Y. Imaging transcription elongation dynamics by new technologies unveils the organization of initiation and elongation in transcription factories. Current Opinion in Cell Biology 74, 71–79 (2022).

107 Sabari, B. R. et al. Coactivator condensation at super-enhancers links phase separation and gene control. Science 361, eaar3958 (2018).

108 Cho, W.-K. et al. Mediator and RNA polymerase II clusters associate in transcription-dependent condensates. Science 361, 412–415 (2018).

109 Donovan, B. T. & Larson, D. R. Regulating gene expression through control of transcription factor multivalent interactions. Molecular Cell 82, 1974–1975 (2022).

110 Oren, M. Regulation of the p53 tumor suppressor protein. Journal of Biological Chemistry 274, 36031–36034 (1999).

111 Harris, S. L. & Levine, A. J. The p53 pathway: positive and negative feedback loops. Oncogene 24, 2899–2908 (2005).

112 Navalkar, A. et al. Oncogenic gain of function due to p53 amyloids occurs through aberrant alteration of cell cycle and proliferation. Journal of cell science 135, jcs259500 (2022).

113 Alberti, S. & Hyman, A. A. Are aberrant phase transitions a driver of cellular aging? BioEssays 38, 959–968 (2016).

114 Dai, Z., Li, G., Chen, Q. & Yang, X. Ser392 phosphorylation modulated a switch between p53 and transcriptional condensates. Biochimica et Biophysica Acta (BBA)-Gene Regulatory Mechanisms 1865, 194827 (2022).

115 Bouchard, J. J. et al. Cancer mutations of the tumor suppressor SPOP disrupt the formation of active, phase-separated compartments. Molecular cell 72, 19–36 (2018).

116 Sabari, B. R. et al. Coactivator condensation at super-enhancers links phase separation and gene control. Science 361, eaar3958 (2018).

117 Chandra, B. et al. Phase separation mediates NUP98 fusion oncoprotein leukemic transformation. Cancer discovery 12, 1152–1169 (2022).

118 Zuo, L. et al. Loci-specific phase separation of FET fusion oncoproteins promotes gene transcription. Nature communications 12, 1491 (2021).

119 Powers, S. K. et al. Nucleo-cytoplasmic partitioning of ARF proteins controls auxin responses in Arabidopsis thaliana. Molecular cell 76, 177–190 (2019).

120 Patel, A. et al. A liquid-to-solid phase transition of the ALS protein FUS accelerated by disease mutation. Cell 162, 1066–1077 (2015).

121 Xiao, R. et al. Pervasive chromatin-RNA binding protein interactions enable RNA-based regulation of transcription. Cell 178, 107–121 (2019).

122 Oksuz, O. et al. Transcription factors interact with RNA to regulate genes. Molecular Cell 83, 2449–2463 (2023).

123 Riback, J. A. et al. Compositional-dependent thermodynamics of intracellular phase separation. Nature 581, 209–214 (2020).

124 Elbaum-Garfinkle, S. et al. The disordered P granule protein LAF-1 drives phase separation into droplets with tunable viscosity and dynamics. Proceedings of the National Academy of Sciences of the U.S.A. 112, 7189–7194 (2015).

125 Weber, S. C. & Brangwynne, C. P. Getting RNA and protein in phase. Cell 149, 1188–1191 (2012).

126 Caudron-Herger, M. & Rippe, K. Nuclear architecture by RNA. Current opinion in genetics & development 22, 179–187 (2012).

127 Zhang, H. et al. RNA Controls PolyQ Protein Phase Transitions. Molecular Cell 60, 220–230 (2015).

128 Adame-Arana, O., Bajpai, G., Lorber, D., Volk, T. & Safran, S. Regulation of chromatin microphase separation by binding of protein complexes. Elife 12, e82983 (2023).

129 Shin, Y. et al. Liquid nuclear condensates mechanically sense and restructure the genome. Cell 175, 1481–1491 (2018).

130 Jambon-Puillet, E. et al. Phase-Separated Droplets Swim to Their Dissolution. bioRxiv, 10.1101/2023.07.18.549556 (2023).

