## Supplementary Information for "Nucleo-cytoplasmic environment modulates spatio-temporal p53 phase separation"

**Contents** **Page no.**


| | |
|---|---|
| 1. Materials and Methods | 3-23 |
| 2. Supplementary Figures (1-15) | 24-38 |
| 3. Supplementary Table (1-3) | 39-40 |
| 4. Supplementary movie legends (1-6) | 41 |
| 5. References (1-23) | 42-43 |

### MATERIALS AND METHODS

#### Chemicals and reagents

The chemicals and reagents used for the experimental purposes were majorly procured from Sigma Aldrich Co (St. Louis, MO, USA) or Merck (Darmstadt, Germany) unless mentioned otherwise. The water was double distilled and deionised by a Milli-Q water purification system (Millipore Corp., USA).

#### *In silico* analysis of p53 protein

Full-length p53 and the p53 core domain (p53C) sequence were used for *in silico* analysis. The disordered regions of the primary structure of the protein were identified using the Online tool IUPred2A[1,2] and the droplet-promoting region was predicted using Fuzpred[3]. SMART[4] (Simple Modular Architecture Research Tool) was used to identify low complexity regions (LCR) along the protein sequence. Output data was plotted using OriginPro 2021 (Origin Lab, USA) software. The secondary structure of p53 and p53C was obtained from Alphafold[5] and Protein Data Bank[6] (PDB ID:6OCJ). The output PDB structures were represented using PyMOL[7].

#### Site-directed mutagenesis

Plasmids for expression of GFP-p53 WT, GFP-p53 NES- and p53C WT [p53C WT (residues 94-312)] were obtained from Addgene (Addgene ID # 12091, 12092 and 24866, respectively). The mammalian and bacterial expression constructs for p53 mutants were constructed via Polymerase Chain Reaction (PCR) mediated site-directed mutagenesis. The plasmid containing GFP-p53 WT was used as the template for two mammalian expression constructs of p53 mutants GFP-p53 R175H and GFP-p53 R248Q. For bacterial expression constructs containing mutant p53C, namely, p53C R175H and p53C R248Q, p53C WT (residues 94-312) was used as the template. Primers were designed against the template by altering the codon at the mutation site such that the mutation site lies at the primer's center (**Table S1**). Pfu DNA

Polymerase (SRL Ltd, India) was used for polymerase chain reaction (PCR). After 20 thermal cycles, the PCR product was selected via Dpn1 (Invitrogen, USA) digestion. The digested PCR product transformed XL10 (Gold) competent cells, and colonies containing the construct were selected using relevant antibiotic resistance (Kanamycin for GFP-p53 variants and Ampicillin for p53C variants) in Luria-Agar (LA) medium. The colonies were thereby grown on Luria broth (LB) media. Plasmids were isolated by mini-prep kit (Favorgen Biotech Corp., Taiwan) and quantified using a NanoDrop UV photometer (Implen, USA). The newly generated plasmids were verified by sequencing analysis.

**Cell lines and transfection studies**

MDA-MB-231 (human breast epithelial adenocarcinoma) cells, MCF7 (human breast epithelial adenocarcinoma) cells, HeLa cells (human cervical epithelial adenocarcinoma), MCF 10A (human mammary gland epithelial cells) and SaOS2 (human bone epithelial osteosarcoma) cells were obtained from the National Centre for Cell Science (NCCS, Pune, India). MCF 10A, MDA-MB-231 and MCF7 cells were seeded on coverslips, allowed to grow until 80% confluency, and fixed for immunostaining. For transfection studies, cells were seeded on 35 mm confocal dishes and allowed to grow to 70% (HeLa) and 90% (SaOS2) confluency. Hereafter, the cells were transfected with 1 μg of DNA using Lipofectamine and P3000 following the manufacturer's protocol (L3000-008, Invitrogen, USA). HeLa cells were cultured in DMEM containing 10% fetal bovine serum (FBS, Gibco, USA), while SaOS2 cells were cultured in McCoy's 5A Medium (Cat# M4892, Sigma Aldrich) with 15% FBS. The cells were maintained at 37$^{o}$C with 5% $CO_2$. For microscopic studies, Cisplatin (Cat# P4394, Sigma Aldrich) and Leptomycin B (Cat# L2913, Sigma Aldrich, USA (20 ng/mL) were added after 12 h post transfection at concentrations 10 μM and 20 ng/mL, respectively.

**Immunostaining of cells**

MCF 10A, MCF7 and MDA-MB-231 cells were first seeded on 12 mm coverslips. At 80% confluency, cells were fixed with 4% paraformaldehyde solution for 20 min at 37 $^{0}$C. Fixed cells were hereby permeabilised using 0.2% Triton X in Phosphate buffer saline (PBS) for 15 min, and nonspecific antigen sites were blocked with 2% BSA for another 1 h. Cells were then stained with mouse monoclonal anti-human p53 (DO-1) (Cat# sc-126, Santa Cruz Biotechnology, USA) antibody (1:200) overnight at 4 $^{0}$C. The coverslips were then washed with PBST (0.1% Tween 20 in PBS). The coverslips were incubated with Alexa Fluor 555 (Cat# A32727, Invitrogen, USA) or Alexa Fluor 488 (Cat# A11001, Invitrogen, USA) conjugated goat anti-mouse secondary antibody diluted in PBST (1:500) for 2 h at 37 $^{0}$C for MDA-MB-231 and MCF 10A cells and (1:250) for 3 h at 37°C for MCF7 cells. Cells were washed with PBST and 4',6-diamidino-2-phenylindole (DAPI) (1 µg/mL) was added for 2 min for nuclear stain. Cells were then washed with PBS and mounted on a glass slide with mounting medium containing 1% (w/v) 1,4-Diazabicyclo [2.2.2] octane (DABCO) and 90% (v/v) glycerol in PBS. Images were acquired using Laser Scanning confocal microscope (Zeiss LSM 780, Carl Zeiss, Germany) with iPlan-apochromat 100x/1.4 NA objective.

For SaOS2, cells were seeded on 12 mm coverslips and transfected with GFP-p53 WT, R175H and R248Q and fixed at early (18 h) and late (48 h) time points with ice-cold 4% paraformaldehyde (PFA) in PBS for 5 min at 37°C for immunostaining of misfolded p53. Using similar protocol as mentioned before, cells were stained with mouse monoclonal anti-human Pab240 antibody (1:500) (Cat# sc-99, Santa Cruz Biotechnology, USA) followed by Alexa Fluor 555 conjugated anti-mouse secondary antibody (1:500). Images were acquired using Spinning Disk confocal microscope (Zeiss Axio-Observer Z1 with Nipkow spinning disc), using iPlan-apochromat 63x/1.4 NA objective. SaOS2 cells were transfected with GFP-p53 WT as described previously. Post 18 h and 48 h transfection, cells were stained with

MitoTracker Red (Cat# M7513, ThermoFisher Scientific, USA) and LysoTracker Red DND-99 (Cat# L7528, ThermoFisher Scientific) as per the manufacturer's protocol. For the Nile red staining, Nile red (10 μg/mL) was added to cells at 18 h and 48 h post transfection and incubated for 3 min. The cells were washed using PBS after incubation and Opti-MEM (Cat# 11058021, Gibco) was added before capturing the images. For ProteoStat binding assay, 0.5 μL of ProteoStat solution (5 μM) was diluted in 1 mL of assay buffer provided in the kit (Enzo Life sciences, USA). The solution was added to the fixed cells (at 18 h and 48 h post transfection) and incubated for 30 mins at RT as per manufacturer's protocol. The cells were washed using PBS and Opti-MEM was added before imaging. All the images were acquired using Zeiss Laser scanning confocal microscope (LSM 780, Carl Zeiss, Germany) with iPlan-apochromat 100X/1.4 oil objective. The images were processed using FIJI software.

**Time-lapse confocal microscopy imaging**

Live cell imaging was done for time-dependent studies post-transfection of GFP-p53 with HeLa and SaOS2 cells. The time-dependent cellular imaging was done at 12 h, 18 h, 24 h, 36 h and 48 h (512 x 512, 8 bit) and images were captured using Spinning Disk confocal microscope (Zeiss Axio-Observer Z1 with Nipkow spinning disc) using 63X/1.4 NA (oil) or 100X/1.4 NA (oil) objectives. Opti-MEM (Cat# 11058021, Gibco) was used as media for all live cell imaging and incubation was maintained at 5% $CO_2$ and 37 °C. For nuclear stain, Hoechst dye (Cat# 33342, Enzo, USA) was used. For *in vitro* characterisation of p53C LLPS, NHS-Rhodamine labelled p53C was used for imaging at 37 °C. The excitation source of 488 nm solid-state laser was used for GFP-p53 and Atto 488 conjugated DNA and 561 nm solid-state laser was used for NHS-Rhodamine labelled p53C. Images and movies were processed using Zen Blue software (Zen Lite 2012).

**Western blotting**

At various time points post transfection, HeLa and SaOS2 cells were suspended in 1X Laemmli sample buffer (62.5 mM Tris pH 6.8, 2% SDS, 10% Glycerol) for protein extraction. The suspension was centrifuged at 20817 X g at 20 °C for 1 h, and the supernatant was collected for protein estimation using BCA kit (Pierce BCA Protein Assay Kit, Cat# 23227, Thermo Fischer Scientific, USA). Further, 40 µg of protein was loaded for separation by SDS-PAGE electrophoresis and blotted onto a 0.22 µm PVDF membrane (Cat# BSP0161, Pall life sciences). After transfer, the PVDF membrane was blocked with 5% BSA in TBST for 1 h at room temperature. The blot was probed with the respective primary antibody made in 2% BSA in TBST with 1:2000 dilution and incubated overnight at 4 ºC. The PVDF membrane was washed with TBST and the blot was incubated with the HRP-conjugated secondary antibody (1:2000 dilution prepared in 2% BSA in TBST) and incubated at room temperature for 1 h. The PVDF membrane was then washed with TBST and signals were detected using Clarity Western ECL detection kit (Cat# 1705060, Bio-Rad) according to the manufacturer's instructions. The images were captured using the Image Quant LAS 500 (GE life sciences, USA). The captured images were quantified using the Biorad Image lab software. The antibodies used for western blotting experiments are anti-GAPDH (Cat# sc-365062, Santa Cruz Biotechnology, USA), anti-p53 (Cat# MA5-16387, Invitrogen, USA), anti-p21 (Cat# 2947, Cell Signalling Technology, USA), anti-histone H3 (D1H2) (Cat# 4499, Cell Signalling). The secondary antibodies used were Goat anti-mouse IgG, H & L Chain Specific Peroxidase Conjugate (Cat# 401253, Merck, USA) and goat anti-rabbit IgG, H & L Chain Specific Peroxidase Conjugate (Cat# 401353, Merck, USA).

**Fluorescence recovery after photobleaching (FRAP) experiments**

FRAP studies were done using Laser Scanning Confocal microscope (Zeiss LSM 780 Axio-Observer Z1 microscope). HeLa cells and SaOS2 were seeded on 30 mm glass bottom confocal dishes (Genetix, India) and transfected with plasmids containing mammalian expression vectors with GFP-p53 WT, R175H and R248Q, separately. At various time points, the dish was mounted on the microscope with incubation temperature 37 °C, 5% $CO_2$ and under optimal humidity conditions during imaging. At each time point (18 h and 48 h), photobleaching of nuclear and cytoplasmic condensates was done using 488 nm laser (100% laser power). Along with the region of interest (ROI) for photobleaching, 2 more ROIs were selected within and outside condensates, respectively, for bleaching correction. Time-lapse images were taken with laser excitation at 488 nm and emission was recorded at 534 nm. Recovery was monitored until there was no signal strength change with time, indicating the limit of recovery of the bleached region. For FRAP studies of nuclear condensates, due to small size and high dynamicity, translational correction of the images was done post-acquisition using FIJI linear stack alignment tool[8]. For this, new ROIs were selected in the aligned, and the intensity of the whole stack for each ROI was recorded and processed to calculate fluorescence recovery in a similar method as stated before.

For FRAP studies with p53C LLPS *in vitro*, 10% labelled protein [10% (v/v) NHS-Rhodamine labelled protein mixed with 90% unlabelled protein] was used. Protein labelling was done as per previously published protocols[9]. At different time intervals, photobleaching of the droplets was done using 561 nm DPSS 561-10 laser (100% laser power). For every bleaching, 2 unbleached ROIs of the same diameter inside and outside of the droplets were taken as controls for bleaching correction. Post bleaching, the recovery was monitored until a plateau in the signal was reached. Objectives iPlan-apochromat 100X/1.40 (oil) with GaAsP detector was used for cytoplasmic, nuclear and *in vitro* p53C condensates. The intensity profiles were recorded using Zen Pro (Zeiss, Germany) software with 8-bit depth. For both *in cellulo* and *in vitro* FRAP studies, the data was normalised after bleach correction and fitted in Origin Pro 2021 (Origin Lab, USA) software using exponential growth function for calculation of half-time ($t_{1/2}$) as per protocol mentioned in Ray *et al* [10].

**STED imaging of p53 condensates**

Super-resolution studies of p53 nuclear and cytoplasmic condensates were done using stimulated emission depletion (STED) microscopy[11,12] (Leica Microsystems, Germany). For cells, MDA-MB-231 and MCF7 cells were seeded, fixed on coverslips, and immunostained as described in the previous section. Post staining, the cells were mounted on glass slides with Mowiol mounting medium [10% (w/v) Mowiol 4-88, 25% (v/v) Glycerol in 100 mM Tris-HCl (pH 8.5)]. For GFP-p53 variants, SaOS2 cells were seeded and transfected with GFP-p53 WT, R175H and R148Q plasmids and fixed at early (18 h) and late (48 h) time points with ice-cold 4% paraformaldehyde in PBS for 5 mins. For stressor treatment, 10 µM of Cisplatin (Cat# P4394, Sigma Aldrich) was added at an early time point (18 h) and fixed after 3 h. For all the cells, coverslips containing cells were mounted on a glass slide with Mowiol mounting medium. Images were captured with STED microscope (Leica Microsystems, Germany) and post-acquisition, images were deconvoluted using CMLE algorithm[13] in Huygens Professional Suite. Rendering of microscopic images for visualisation of condensates was carried out using IMARIS 8.3.1 (Bitplane, Switzerland). For quantification of the percentage of p53 condensate occupancy with the total pool of p53 in the nucleus, the thresholding tool[14,15] of FIJI software was used. The whole area of the nucleus was first selected before thresholding to limit the area of estimation. The intensity thresholding tool of p53 was used to estimate the total p53 in the nuclear region, and the local thresholding tool[16,17] was used to identify the portion of nuclear condensates that had higher local intensity relative to its surroundings due to clustering. The percentage of nuclear occupancy was calculated as % = (area of the region above local threshold/area of the region above threshold) X 100. Trainable Weka Segmentation tool[18] of FIJI was used for estimating the area of the nuclear condensates. Each image was trained using 50 regions each for condensate (region 1) and 50 for background (region 2). The segmentation classifier was then run and the resultant segmented image was obtained, defining the clusters and background. Thresholding was done on this segmented image to obtain the area and Feret diameter was calculated for each condensate. For the high irregularity of the shape of the nuclear clusters, the area was taken as a measure of the size of the condensates. Considering the lowest possible resolution of images obtained by STED to be 50 nm, areas above 2500 $nm^2$ were counted for estimation. The area distribution plots for each pair for each variant (GFP-p53 WT, R175H and R248Q), were pairwise normalised before plotting to account for the different sizes of the nucleus. The data was plotted using MATLAB (MathWorks) 2019a software.

**Lattice light-sheet microscopy of p53 condensates.**

Lattice light-sheet microscopy (LLSM) was done to examine nuclear condensates in live SaOS2 cells. For this, the cells were seeded on 5 mm coverslips and allowed to grow to ~80% confluency. Hereafter, cells were transfected with WT GFP-p53 plasmid and imaged at 18 h and 48 h post transfection using 3i Lattice light-sheet microscope (3i, Colorado, USA). Volumetric image stacks were generated using a square lattice in dithered mode. A square lattice was made for the 488 nm channel with 51 beams spaced at 0.99 µm intervals between the beams, and the pattern was cropped with a cropping factor of 0.150. The deskewed step size was 0.163 um after scanning the sample stage in 0.3 µm steps. The excitation laser power was adjusted between 0.14 mW and 0.43 mW at the image plane to obtain a good signal to noise ratio. In the emission path, a custom-made 488-640T/560 R dichroic beam splitter was used. A quad-notch filter (Semrock FF01-446/523/600/677) was added to the transmitted path to collect the emitted signal. The final fluorescence signal was imaged on a Hamamatsu ORCA-

Fusion Bt sCMOS camera. Each image plane was captured with a 10 ms exposure time. The frame size of the image in X, Y, and Z directions was between 20-50 µm, 20 µm, and 140- 200 planes (23-30 µm), respectively to ensure the entire nuclear region was imaged while avoiding any strong fluorescence signal near the edges of the frame. Hence, a volumetric image stack of 50x20x25 µm was captured within ~ 2.5 s (100 planes for 1.2 s).

Post acquisition, the images were cropped out from the raw datasets. Since the images were captured at an angle of approximately 32 degrees using LLSM, the images were deskewed and corrected for coverslip orientation in the Slidebook software. Photobleach correction was applied when necessary. The images were further deconvoluted using Richardson Lucy Constraint Iterative (CI) algorithm with 10 iterations using theoretical point spread function and a Gaussian noise smoothening radius of 0.3. Final images were processed in FIJI for representative purposes.

**Apoptosis studies using flow cytometry**

For studying the time-dependent (6 h, 12 h, 18 h, 24 h, 36 h and 48 h) apoptotic cell death upon WT p53 transfection cells, SaOS2 (7 x $10^5$ cells/well) were seeded in six-well plates (Corning, USA). After 24 h, cells were transfected with GFP-p53 WT plasmid in a time-dependent manner, i.e. 48 h, 36 h, 24 h, 18 h, 12 h and 6 h before the acquisition using flow cytometry. The cells were trypsinised and collected by centrifugation at 664 X g for 4 min at 25°C. The cell pellets were washed with ice-cold PBS and centrifuged at 664 X g. The cell pellets were suspended in 100 µL of 1X binding buffer followed by doubling staining with V450 Annexin V and PI (Cat# 560506, BD Horizon) and incubated for 15-20 min at room temperature. The samples were diluted to 400 µL with the 1X binding buffer and quantified by flow cytometry (FACS Aria, BD Biosciences). The gating was performed using untransfected cells and ~10,000 GFP-p53 WT positive cells were counted for each sample. The values are normalized using untransfected control cells. The recorded data were plotted using FlowJo v10 software (Tree Star, Inc).

##### Chromatin immunoprecipitation (ChIP)-qPCR assay

ChIP-qPCR assay was performed to determine the functional status of p53 protein. For this, SaOS2 cells were transfected with WT p53 and mutant p53 (R175H and R248Q) as per the described protocol[19]. The cells were collected at 18 h and chromatin was cross-linked using 1% formaldehyde for 30 min and added directly to the media. Further, the cross-linking was quenched using 125 mM glycine for 5 min. To attain higher degree of resolution during detection, the samples were sonicated to obtain DNA fragments that are ~200 -1000 bps in size. Sonication was done at 30% amplitude for a total of 3 cycles with 1 min ON and 1 min OFF on ice. The samples were centrifuged at 10,000 X g at 4 °C for 10 min to remove insoluble material. The cells were incubated overnight using 5 µg (25 µl) of p53 antibody (DO-1) as described earlier (25). The chromatin-bound DNA was eluted with 50 µl of elution buffer using Magna ChIP™ G Tissue Kit (Millipore, USA) as per the manufacturer's instructions. QIAquick PCR purification kit (QIAGEN, Valencia, USA) was used for purification of all DNA samples (IPs and inputs) in accordance to the manufacturer’s protocol. Further, qPCR was performed with SYBR Green reaction mixture (20 µl) using Agilent qPCR machine. Both ChIP and qPCR experiments were performed in duplicates. Calculations for enrichment/input values were made using the equation 1,

$$\Delta \mathrm{CT} = \mathrm{CT(ChIP)} - [\mathrm{CT(Input)} - \mathrm{LogE}\ (\text{Input dilution factor})]^{-\Delta CT} \qquad (1)$$

where E represents the specific primer efficiency value and % Enrichment/Input=E. The sequence of primers for p53 response element for p21 was as follows:

Forward primer-5’-GTGGCTCTGATTGGCTTTCTG-3’

Reverse primer: 5’- CTGAAAACAGGCAGCCCAAG-3’

**p53 DNA binding assay in cells**

Enzyme-linked immunosorbent assay (ELISA) for p53 was performed to determine the ability of p53 to bind to the DNA element.For this, SaOS2 cells were transfected with GFP-p53 WT, R175H and R248Q. The cells were collected at 18 h post transfection. The cells were harvested and lysed using RIPA buffer supplemented with Roche protease inhibitor cocktail. The lysate was centrifuged at 4 °C to remove the cell debris. The supernatant obtained was added onto the 96-well plate immobilised with double-stranded DNA sequence containing p53 response element using p53 Transcription Factor Assay kit (Cat# 600020, Cayman Chemicals, Michigan) as per the manufacturer's protocol. p53 present in the cell lysate binds specifically to the p53 response element and is detected by adding a p53 antibody (DO-1). Anti-mouse secondary antibody conjugated to HRP was added to provide a sensitive colorimetric readout at 450 nm. Nutlin-3 stimulated MCF7 nuclear extract was used as a positive control for the assay.

**RNA isolation and qRT-PCR**

SaOS2 cells were seeded on 6-well plates and transfected with GFP-p53 WT. For stressor treatment, 10 μM cisplatin was added at 9 h post transfection. The cells were trypsinised for both treated and untreated cases and pelleted down at 18 h. The RNA isolation was carried out using TRizol reagent (Cat# 15596018, Ambion by Life Technologies, USA) according to the manufacturer's protocol. Briefly, the harvested cell pellets were lysed in 1 mL of Trizol reagent, followed by the addition of 350 μL of chloroform. The tubes were vortexed vigorously and kept in ice for 5 min. The tubes containing lysed samples were centrifuged at 12,000 X g for 15 min at 4 °C. The upper aqueous phase containing RNA was carefully transferred into new tubes and added half a volume of isopropyl alcohol for the precipitation of RNA. The samples were mixed and incubated for 30 min at room temperature and centrifuged at 12,000 X g for

10 min at 4 °C. The white gel-like RNA pellet was collected and washed with 1 mL of ethanol (70%) and centrifuged at 7500 X g for 5 min at 4 °C. The supernatant was discarded and the RNA pellet was allowed to air dry and resuspended in nuclease-free water. The samples were digested with DNase I to remove the DNA contamination from the isolated RNA sample as per the manufacturer's instruction (Turbo DNA-free kit, Cat# AM-1907, Thermo Fischer Scientific, USA). The concentration of the purified RNA was measured using a nanodrop spectrophotometer (Implen, USA). Hereafter, cDNA was synthesised from total RNA using RevertAid First Strand cDNA Synthesis Kit (Cat# K1622, Molecular biology, Thermo Scientific, USA) as per the manufacturer's instruction. Oligo (dT) primers were used for the reverse transcriptase reaction. Real-time PCR (qRT-PCR) was further carried out on a CFX96 Touch Real-Time PCR detection system (Bio-Rad, USA) using the SYBR Green method. Maxima SYBR Green/ROX qPCR Master Mix (2X) (Cat# K0221, Thermo Scientific, USA) was used according to the manufacturer's protocol with the predesigned primers for the required genes (**Table S3).** Two independent experiments in duplicates were performed for each reaction.

**p53 core (p53C) protein expression and purification**

Plasmid (pet15b vector carrying 6XHis tagged 94-312 amino acids gene sequence) for p53C (WT, R175H and R248Q) was transformed into BL21(DE3) competent cells using the standard protocol. Transformed cells were selected based on Ampicillin resistance. A single colony was inoculated in 100 mL Luria Broth (LB) and grown to $OD_{600}$ of 0.5 as a starter culture. Hereafter, 20 mL of starter culture was added to 1 L LB with antibiotic and allowed to grow till $OD_{600}$ of 0.7. The protein expression of WT, R175H and R248Q were induced by 1 mM, 0.25 mM and 1 mM isopropyl β-D-1-thiogalactopyranoside (IPTG), respectively, for 12 h at 18 °C. The cells were then harvested and resuspended in lysis buffer (50 mM Sodium phosphate, 300 mM Sodium chloride). 1X protease inhibitor cocktail (Cat# 05056489001, Roche cOmplete, Mini,

EDTA-free, Roche Diagnostics, Germany) was added to prevent proteolysis. The cell suspension was then sonicated using a probe sonicator (Sonics & Materials, Inc, pulse of 2 sec ON, 2 sec OFF; 40% Amplitude per cycle, for 5 cycles). The soluble fraction was collected and passed through the column of the Qiagen Nickel NTA (Cat# 1018244, Qiagen, Germany). The column was washed with lysis buffer containing 50 mM Imidazole (4 column volumes) and protein fractions were eluted against gradient concentration of imidazole from 100 mM to 500 mM. Each gradient was run on SDS-PAGE to identify the fraction containing the maximum yield of the protein. Hereafter, the protein was passed through size exclusion chromatography (SEC) (Hi Load 16/60, Superdex 200 TM 10/300, Cat# 17-1069-01, Cytiva Life sciences) column pre-equilibrated with 2 column volumes of 50 mM sodium phosphate buffer (PB) (pH 7.4, 0.01% sodium azide) for further purification. The eluent protein was then further used for labelling and experimental purposes.

The protein concentration was measured using UV spectroscopy following Beer Lambert's law. The absorbance was recorded at 280 nm and concentration was estimated considering the molar absorptivity constant as 17420 $M^{-1}cm^{-1}$.

For experimental studies involving labelled p53C, the protein was labelled with NHS-Rhodamine (Cat# 46406, Thermo Scientific, USA), using the manufacturer's instructions in 50 mM PB. A five-fold molar excess of NHS-Rhodamine was added to p53C and incubated at 4 °C for 3 h with constant stirring. Hereafter, the excess dye in the labelled protein mixture was then dialysed out using 12.4 KDa cut-off membrane (Cat# D0655, Sigma Aldrich, USA) in 50 mM PB. The efficiency of labelling was verified by calculating the degree of labelling before proceeding with the experiment as per manufacturer protocol. The concentration of the labelled protein was determined using spectroscopic methods as per instructions.

##### *In vitro* p53C phase separation studies

SEC purified WT, R175H and R248Q p53C were studied for phase separation at varied concentrations in the presence of various concentrations of molecular crowder Polyethylene Glycol (PEG) 8000 (0%, 5%, 10%, 15% and 20%) in 50 mM PB, pH 7.4 to construct the phase regime. For microscopy-based studies, 10% (v/v) NHS-Rhodamine labelled protein was used. Different concentrations of protein in the presence and absence of PEG 8000 were prepared in 50 mM PB (pH 7.4). Glass coverslips (No.1) and glass slides were thoroughly washed with 1% Hellmanex solution, dried, and 10 µl of the reaction mixture was spotted on the coverslips. The coverslips were mounted on the glass slides and were sealed properly using transparent nail paint and incubated at 37 °C in a hydrated chamber for time-dependent studies. Hereafter, at regular time intervals, the samples were visualised for condensate formation under the Zeiss Spinning disk confocal microscope using 63X/1.40 oil objective. For p53C studies along with non-labelled DNA, Atto labelled target (TR) and non-target (NTR) DNA (Sigma Aldrich, USA) was used. 10% (v/v) labelled DNA was used in all the experiments. The imaging of LLPS of p53C samples in the presence of unlabelled DNA and RNA, was captured using Zeiss Laser scanning confocal microscope under 63X/1.40 oil or 100X/1.40 oil objective. Quantification of size and number of condensates were done using the Thresholding tool[16] of FIJI and plotted using KaleidaGraph (v 4.03) and GraphPad Prism 8 software.

##### Light scattering studies

Light scattering studies were performed for 10 µM of p53C in 50 mM PB (pH 7.4) in the presence of PEG-8000 (10% w/v) as crowder at 37 °C. The experiments were performed in continuous mode, with a 1 min time interval using JASCO FP8500 (USA) spectrofluorimeter, with excitation and emission wavelength of 350 nm and 5 nm slit width. 120 µL of p53C WT, R175H and R248Q samples were loaded into quartz cuvette and scattering was recorded immediately. Each experiment was performed twice. For studying the effect of DNA on the LLPS of p53C, 10 μM TR-DNA, 10 μM NTR-DNA and ~20 nM (30 μg/mL) RNA were added to 10 μM of p53C in 50 mM PB, pH 7.4, in the presence of 10% (w/v) PEG-8000 and scattering profile was recorded. Similarly, to study the effect of DNA and RNA on pre-formed p53C condensates, first, scattering reactions were set up for WT p53C in the presence of 10% (w/v) PEG-8000. Hereafter, immediately on attaining saturation in the scattering profile, TR-DNA, NTR-DNA and RNA were added to a final concentration of 10 μM, 10 μM and 30 μg/mL, respectively, and the scattering profile was further recorded post addition in the same settings. The data was plotted using Origin Pro 8 software (Origin Lab, USA).

**Fourier Transform Infrared (FTIR) Spectroscopy**

FTIR spectroscopic studies were done to elucidate the secondary structure of the WT p53C in the dilute and dense phase after phase separation. Initially, 500 μL of 20 μM of protein was taken and that was allowed to phase separate in 50 mM PB in the presence of 10% PEG-8000 for 1 h. The formation of condensates was verified after observation under the microscope. Hereafter, the protein sample was centrifuged at high speed (20,000 X g) for separation of the dilute phase and dense phase[20]. 5 μL of dilute phase (supernatant) and dense phase (pellet) were spotted on separate thin KBr pellets and were subjected to dry under an IR lamp. Then, the spectrum was obtained in an amide I stretching frequency in a range of 1800-1500 $cm^{-1}$, with a resolution of 4 $cm^{-1}$, by using Bruker VERTEX 80 spectrometer (Bruker, Leipzig, Germany) attached with a DTGS detector. The recorded spectrum was baseline corrected and then deconvoluted using Fourier Self Deconvolution (FSD) method at the frequency range of 1700–1600 $cm^{-1}$. The Lorentzian curve fitting method was used to fit the spectrum using OPUS-65 software (Bruker, Germany) according to the manufacturer's instructions. Curve fitting and the area under the curve were determined and the frequencies were noted. Three independent experiments were performed for each sample. The data was plotted using KaleidaGraph (v 4.03) software.

**Transmission electron microscopy**

10 μM of WT, R175H and R248Q p53C samples in 50 mM PB (pH 7.4, in the presence of 10% PEG-8000) were set up for the formation of condensates on coverslips mounted on glass slides. After the formation of condensates (confirmed via observation under the microscope), the samples from the coverslip were transferred to copper formvar EM grid (Electron Microscopy Sciences, USA) and incubated for 5 min. The grids were subjected to staining with uranyl formate (1% w/v) for 5 min. The excess liquid on the grid was dried using filter paper and the sample was set to air dry. Prior to imaging, the samples were further dried using an IR lamp. TEM sample preparation and image acquisition were done for condensates formed at 0 h (immediately after formation) and 12 h. Image acquisition was done using 200 KV Transmission electron microscope (JEOL JEM 2100F, Japan) with 10000X magnification. The images were recorded digitally using the Gatan microscopy suite (Gatan, USA).

**Thioflavin T (ThT) and Congo Red (CR) binding assay**

200 μL of 10 μM p53C was set to phase separate in 50 mM PB (pH 7.4) in 10% PEG-8000. 1 mM ThT was prepared in 20 mM Tris-HCl buffer, pH 8.0, containing 0.01% sodium azide. Then, 2 μL of ThT was added to 200 μL of protein sample. Fluorescence was measured with a cuvette (Hellma, volume 500 μL, path length 10 mm) using a Spectrofluorometer (JASCO FB 8500, USA) with excitation wavelength at 450 nm and emission wavelength was recorded from 460-500 nm. A slit width of 5 nm was used for both emission and excitation. For both time points, i.e. at 0 h and 12 h, the fluorescence obtained at 480 nm was plotted as a function of incubation time. α-Synuclein fibril was taken as positive control and freshly purified p53C was taken as negative control.

For CR binding assay, 100 μM CR was dissolved in 50 mM PB containing 10% ethanol. p53C condensates were formed in the presence of 10% PEG-8000. 5 μL CR solution was mixed with 95 μL of LLPS reaction mixture at 0 h and 12 h. For CR binding, absorbance was measured from 300-700 nm. 5 μL of CR with 95μL of 50 mM PB was taken as control. The data was plotted using GraphPad Prism 8 software.

**Effect of DNA on p53C condensate formation**

Microscopy based studies were done to understand the effect of DNA on p53C condensate formation. For this, 10 μL of 100 μM TR-DNA and NTR-DNA were separately spin-coated at the centre of rectangular glass cover glass and 12 mm glass coverslips, using photoresist spinner (PRS-6K, Ducom Ltd.) at 956 X g for 20 s. Cover glass not coated with DNA was taken as control. After drying the sample for 10 mins, the coverslip was mounted on Zeiss Laser Scanning Confocal microscope [LSM 780 Zeiss Axio-Observer Z1 microscope (inverted)] using 100X/1.4 NA oil immersion objective. Additionally, 10 μM of WT p53C was incubated with 10% PEG-8000 in 50 mM PB, pH 7.4, to form condensates, which was confirmed using a microscope. 10 μL of freshly pre-formed condensates were spotted at the centre of the coating on the rectangular coverslip mounted on the microscope. The 12 mm coated coverslips with the same DNA on the corresponding rectangular coverslip were mounted over the sample to avoid evaporation. Pre-formed condensates were also spotted on uncoated cover glass and covered with circular uncoated coverslips, taken as control. For condensates spotted on each case (TR-DNA coated, NTR-DNA coated and uncoated surfaces), images were acquired using Laser Scanning confocal microscope. For each of these conditions, the total number of condensates was calculated using the thresholding tool of FIJI and plotted in GraphPad Prism 8 software.

In another experimental study, we studied the effect of DNA on the extent of partitioning of p53. Here, rubber gaskets (Silicone isolators, Cat# 666505, Grace Bio labs, USA) were mounted over rectangular glass coverslips to create a chamber for holding the LLPS solution. 60 μL of 20 μM protein and PEG-8000 (10% w/v) solution was added to two chambers of the gasket and the solution was allowed to phase separate. Once condensates were formed, as observed under the microscope, images were acquired using Laser Scanning Confocal microscope and the coordinates of the specific positions were marked. Hereafter, TR-DNA and NTR-DNA were added separately to each of the chambers to an equimolar concentration. Images were acquired for the same positions post-addition of DNA, using the same laser settings for the same set of positions. This enabled comparison at a single droplet stage to elucidate the effect of DNA. The extent of partitioning was calculated as the ratio of the intensity of the dense phase to the intensity of the liquid phase using FIJI software. Average fluorescence intensity from ROI of identical size, inside and outside the condensate, was measured, and their ratio was calculated as a measure of the extent of partitioning. The images were acquired in Laser Scanning Confocal microscope using 100X oil objectives.

**Determination of binding affinity of p53 with the TR-DNA, NTR-DNA and RNA**

p53C was labelled using RED-NHS 2$^{nd}$ Generation protein labelling kit (Cat# MO-L011, NanoTemper Technologies GmbH)[21]. Briefly, three-fold molar excess dye was added to 10 μM of p53C and incubated for 30 min at room temperature. The excess unbound dye was removed using the B-column of the labelling kit. The protein concentration was measured using a Nanodrop spectrophotometer (Implen, USA), and the final concentration was diluted to 1 μM in 50 mM PB (pH 7.4). Stock solution of 250 μM of TR-DNA, 250 μM of NTR-DNA was prepared. For determination of binding affinity, serial dilution of ligands (TR-DNA and NTR-DNA) was prepared to a final volume of 10 μL (15625 nM to 0.95 nM for TR-DNA, 50000 nM to 39 nM for NTR-DNA in 50 mM PB, pH 7.4). 10 μL of 200 nM labelled p53, which is the target of the ligand was added to each of these dilutions to a final protein concentration of 100 nM, mixed and incubated for 10 min in dark. The samples were loaded into capillaries (Monolith Premium Capillaries, Cat# MO-K025) and inserted into the Monolith X device (NanoTemper, Germany). The spectral shift measurements at 650 nm and 670 nm were recorded. The experiment was performed three times. The data were normalised, fitted and Kd (binding affinity) was determined using the binding saturation equation (2),

$$y = Bmax * x / (Kd + x) + NS * x + background \qquad (2)$$

where y is the normalised ratio of 670 nm/650 nm, x is ligand concentration, maximum specific binding is denoted by Bmax, equilibrium dissociation constant is represented by Kd, slope of nonlinear regression is represented by NS and background is, measured binding with no added ligand. The data were plotted using the GraphPad Prism 8 software.

**Proteinase K (PK) Digestion Assay**

50 µM of p53C protein sample (without PEG-8000, to avoid the effect of PEG for PK digestion) in 50 mM PB, pH 7.4, was incubated for 12 h at 37 °C to form condensates without any crowder, as confirmed under Fluorescence microscope (DMi8 microscope, Leica Microsystems, Germany). Also, 10 µM of p53C was incubated at identical conditions, as no LLPS control. Hereafter, just prior to the digestion, 50 µM of LLPS sample was quickly diluted to 10 µM in buffer. Both samples (LLPS and no-LLPS) (10 µM) were separately mixed with 10 µg/mL Proteinase K (PK) (Cat# 49936, SRL Pvt. Ltd. India) from a stock of 200 µg/mL. The samples were hereby incubated at 37 °C and the fixed volumes (50 µL) from the reaction mixture were aliquoted at different time intervals (5, 10, 15, 30, 45, 60, 90 and 120 min). The protein samples (LLPS and no-LLPS) just before the addition of PK were taken as samples of 0 min digestion. The reactions at different time points were immediately stopped upon the addition of SDS-PAGE sample buffer followed by heating for 10 min at 95 °C. All the samples

were analysed using 15% Tricine SDS-PAGE (Bio-Rad, USA) and visualised using Coomassie Brilliant Blue stain.

**Differential Scanning Calorimetric (DSC) analysis**

10 μM p53C LLPS and no LLPS samples were prepared the same way as for PK digestion. DSC measurements were run on MicroCal PEAQ-DSC system (Malvern, Worcestershire, UK). The DSC cells were washed with 14% (v/v) Decon 90 solution, followed by a thorough wash with Milli-Q filtered water. First, water measurement was done to verify the performance of the instrument and the cleanliness of the DSC cell. Once repeatable baselines were achieved with water, 50 mM PB, pH 7.4 scan was performed for three times to establish the thermal history of the instrument. Then, three sample scans were examine with the buffer wash in between the sample runs, where freshly prepared samples were loaded for every cycle of heating. For the measurement, 250 μL of 10 μM samples were scanned in the range of 20 °C to 95 °C at a scan rate of 60 °C/h with feedback set to "high", for stability profiling of the protein. Data were analysed using the MicroCal PEAQ-DSC Analysis software (Malvern, Worcestershire, UK). The DSC thermograms of protein samples were corrected for the instrument baseline by subtraction of the corresponding buffer scan. The baseline-subtracted data were fitted to a two-state fitting model to obtain apparent $T_m$ values. The resulting baseline-corrected DSC traces of the p53 protein in LLPS condition and non-LLPS condition were analysed for $T_m$ and the acquired spectra were plotted in Origin Pro 8 (Origin Lab, USA). The experiment was repeated twice with independent sample preparations.

**Separation of nuclear and cytoplasmic fraction of SaOS2 cells**

Nuclear and cytoplasmic fractions of SaOS2 cells were extracted using protocol by Senichkin *et al.*[22,23] Briefly, SaOS2 cells were grown in 60 mm cell culture dishes up to 90% confluency. Hereafter, the cells were washed with PBS, trypsinised and harvested after centrifugation. The pellet-containing cells were washed with PBS once and then resuspended in 200 μL of hypotonic solution [20 mM Tris-HCl (pH 7.4), 10 mM KCl, 2 mM $MgCl_2$, 1mM EGTA, 0.5 mM DTT and 0.5 mM PMSF] and incubated for 5 min at 4 °C. Membrane lysis was done with 0.1% NP-40 for 3 min at 4 °C. The solution was then centrifuged at 1000 X g for 5 min at 4 °C. The supernatant was collected as the cytoplasmic fraction. The pellet containing the nuclear fraction was washed with isotonic buffer [20 mM Tris-HCl (pH 7.4), 150 mM KCl, 2 mM $MgCl_2$, 1mM EGTA, 0.5 mM DTT and 0.5 mM PMSF] containing 0.3% NP-40 in the isotonic buffer for 5 min at 4°C. The solution was centrifuged and the pellet containing nuclei was dissolved in RIPA buffer [25 mM Tris-HCl (pH 7.4), 150 mM NaCl, 0.1% SDS, 0.5% sodium deoxycholate, 1% NP-40, cOmplete™ Protease Inhibitor Cocktail (Roche Diagnostics)] and incubated for 20 min at 4 °C. The solution was again centrifuged at 2000 X g for 3 min at 4 °C and the supernatant was collected as the nuclear fraction. The separated cytoplasmic fraction was centrifuged at 15000 X g to remove the residual debris and supernatant was collected. Post extraction, both the nuclear and the cytoplasmic extracts were dialysed using 1 KDa cut-off membrane (Spectra/Por Dialysis membrane, Cat# 132638, Spectrum Laboratories Inc, USA) in PBS for 2 h. The efficient separation of nuclear and cytoplasmic fractions of SaOS2 cells was further verified by western blotting of the two fractions, following the previously mentioned protocol. An equal protein amount (30 μg), estimated by the Bradford test, was loaded for SDS-PAGE. Anti-GAPDH antibody (1:2000) was used as a cytoplasmic marker and anti-Histone H3 antibody (1:1000, Cat# 4499, Cell Signaling Technology) was used as a nuclear marker. The images were captured using the Image Quant LAS 500 (GE life Sciences, USA).

#### Supplementary Figures

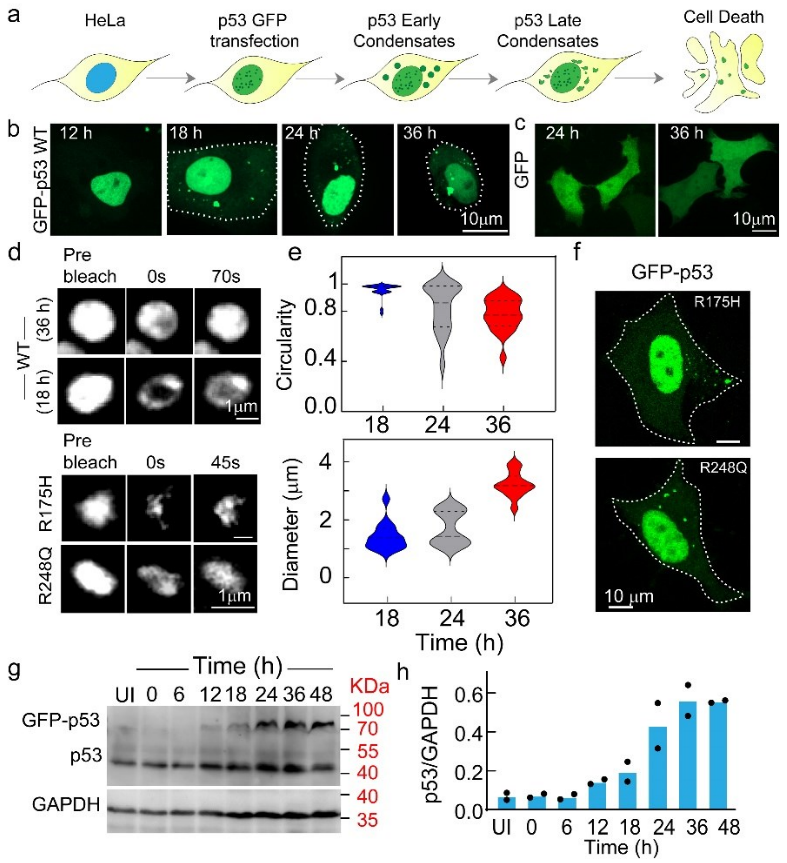


**Figure S1. Transient expression of WT and mutant p53 in HeLa cells. (a)** Schematic representation of transient transfection, expression of p53-GFP and p53 condensate formation in HeLa cells over time. **(b)** Representative confocal images of HeLa cells over-expressing GFP-p53 over time showing p53 condensate formation. The scale bar is 10 μm. The experiment was performed two independent times. **(c)** HeLa cells expressing only GFP showed pan-cellular diffused localization, where no condensates were observed at 24 h and 36 h. The scale bar is 10 μm. The experiment was performed two independent times. **(d)** Representative confocal images of WT p53 ***(upper panel)*** cytoplasmic condensates during FRAP analysis [pre-bleach, bleach (0 s) and post-bleach (the respective time of recovery in second)] at 18 h and late 48 h. FRAP recovery of mutant (R175H and R248Q) ***(lower panel)*** of cytoplasmic condensate at early time point (18 h). The scale bar is 1 μm. The experiment was performed two independent times. **(e)** Violin plots denoting the quantification of the circularity (***upper panel***) and diameter ***(lower panel)*** of p53 WT cytoplasmic condensates in HeLa cells at 18 h, 24 h and 36 h post-transfection. **(f)** Representative confocal images of HeLa cells expressing GFP p53-R175H and GFP-p53 R248Q at 18 h post-transfection showing p53 mutants condensate formation. The scale bar is 10 μm. The experiment was performed two independent times. **(g)** Western blot showing the expression of GFP-p53 (~71 KDa) and endogenous p53 (~48 KDa) with time in Hela cells post-transfection. UI indicating untransfected control. **(h)** The quantification of Western blot showing the expression of GFP-p53 for n=2 independent experiments.

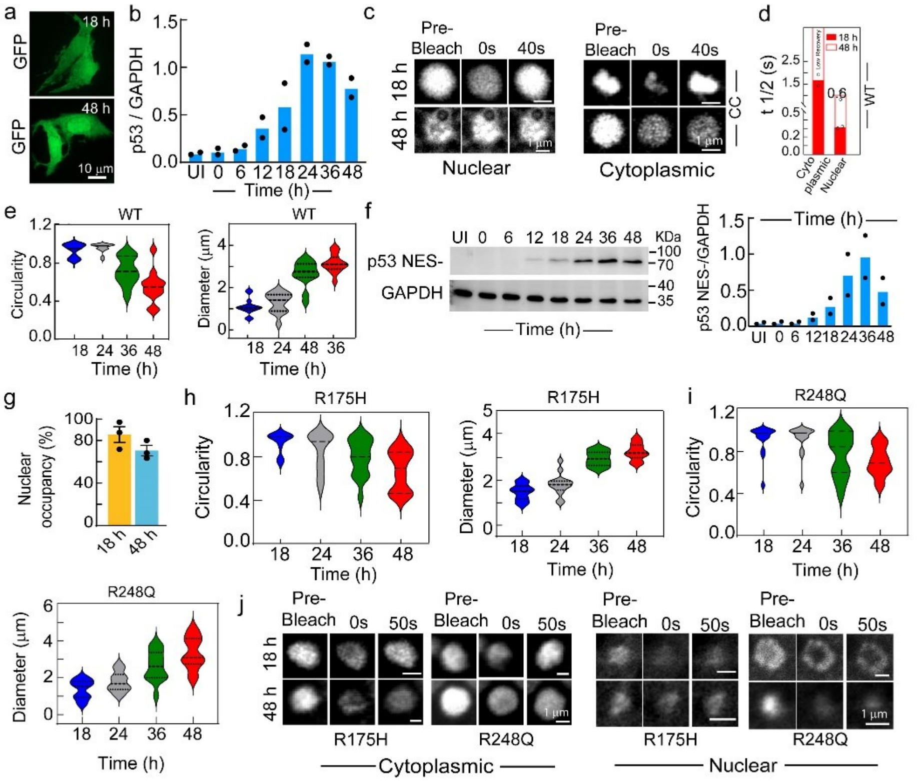


**Figure S2. Characterization of WT and mutant p53 in SaOS2 cells. (a)** The representative confocal microscopy image of SaOS2 cells expressing only GFP showing pan-cellular diffused localization, where no condensates were observed at 18 h and 48 h. The experiment was performed two independent times. The scale bar is 10 µm. **(b)** Fold change of p53 quantified from the Western blot analysis at indicated time points. **(c)** Representative images of p53 cytoplasmic (CC) and nuclear condensates (NC) during FRAP analysis (pre-bleach, bleach (0 s) and post-bleach (40 s) recovery state of condensates) at 18 h and late 48 h. The scale bar is 1 µm. **(d)** Bar graph representing the $t_{1/2}$ post-bleach of early cytoplasmic and nuclear condensate of p53 in SaOS2 cells. Data represents mean for n=2 independent experiments. **(e)** Violin plots showing the quantification of the circularity and diameter of WT cytoplasmic p53 condensate showing a decrease in circularity and an increase in the diameter of the condensates over time. **(f)** Western blot showing the expression of NES- p53 with time. UI indicating untransfected control. ***Lower panel:*** Protein levels were quantified from the Western blot analysis at indicated time points. **(g)** ImageJ analysis of nuclear WT GFP-p53 fluorescence showing p53 condensate as the major population at 18 h and 48 h. The quantification was done from the super-resolution microscopy images. **(h-i)** Violin plots measuring the circularity (***upper panel***) and diameter (***lower panel***) of cytoplasmic condensates of R175H **(h)** and R248Q **(i)** mutant showing an increase in size and decrease in circularity of condensates with time. (**j, k**) Representative confocal images of the cytoplasmic **(j)** and nuclear **(k)** condensates of R175H and R248Q GFP-p53 during FRAP analysis at 18 and 48 h showing pre-bleach, bleach (0 s) and post-bleach states (50 s). The scale bar is 1 µm. The experiment was performed two independent times.

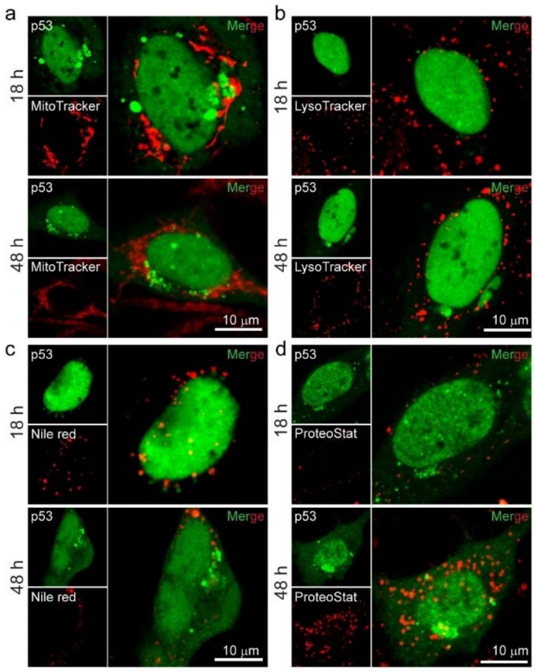


**Figure S3. Characterization of cytoplasmic condensates in SaOS2 cells. (a-d)** Representative confocal images of MitoTracker **(a)**, LysoTracker **(b)**, Nile Red **(c)** and ProteoStat **(d)** staining at 18 h and 48 h post transfection of SaOS2 with WT GFP-p53 showing no colocalization with any of the dye/probe. This suggests the membraneless state of the cytoplasmic condensates are without association of lysosomes, mitochondria and aggreosomes. The scale bar is 10 μm. All the experiments were repeated independently twice with similar results.

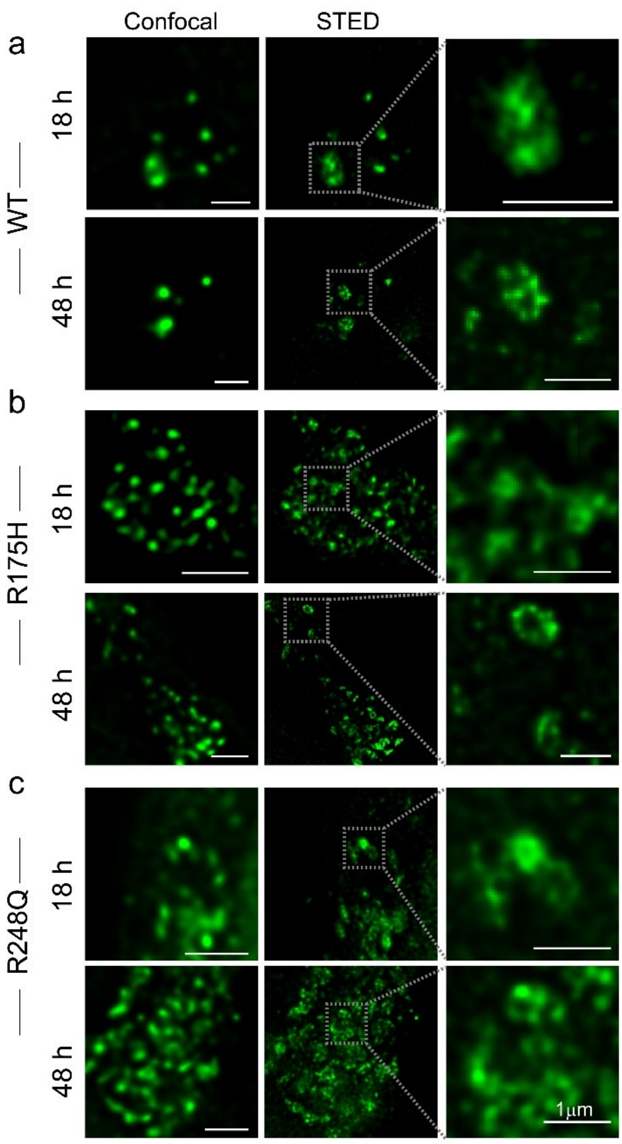


**Figure S4. Super-resolution microscopy (STED) of cytoplasmic condensates by WT p53, R175H and R248Q mutant.** Super-resolution microscopy images of the cytoplasmic condensates of WT **(a)**, R175H **(b)** and R248Q **(c)** p53 at early (18 h) and late (48 h) time points. Cytoplasmic condensates are mostly composed of small condensates. The scale bar is 1 μm. The experiment was repeated twice with similar observations.

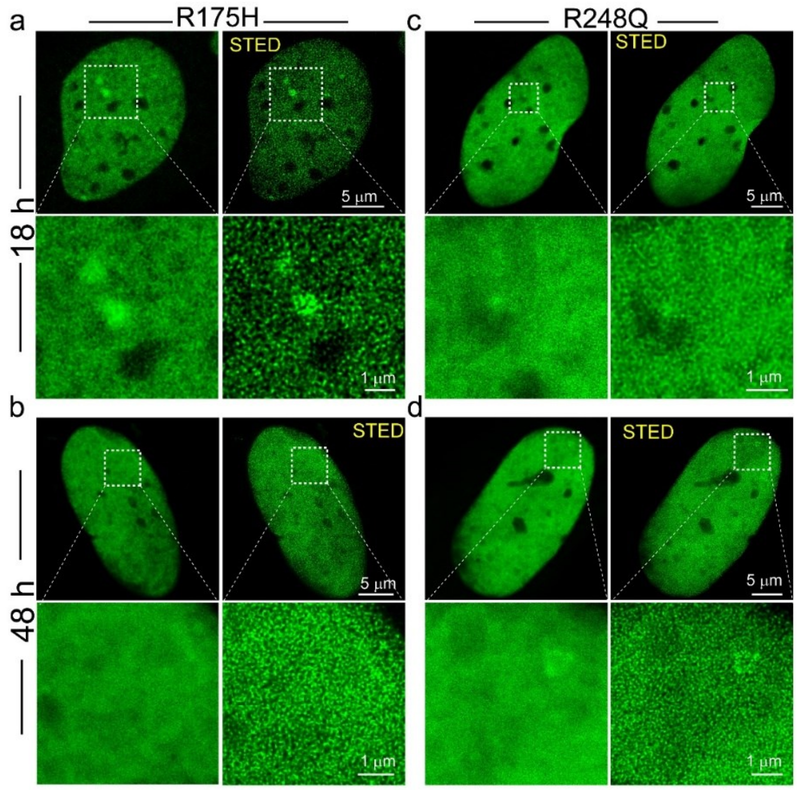


**Figure S5. Confocal and super-resolution microscopy of nuclear p53 mutant (R175H and R248Q) condensates in SaOS2 cells. (a, b)** Confocal microscopy images showing diffused R175H p53 mutant localization ***(left panel***) in SaOS2 cells at early (18 h) and late (48 h) time points. ***Right panel***: Super-resolution microscopy images using STED showing the R175H p53 condensates throughout the nucleus. **(c, d)** Confocal microscopy images showing diffused R248Q p53 mutant localization ***(left panel)*** in SaOS2 cells at early (18 h) and late (48 h) time points. ***Right panel***: Super-resolution microscopy images using STED showing the R248Q p53 condensates throughout the nucleus. The scale bar is 5 µm**.** The zoomed panels represent the magnified area showing a high density of p53 condensates in the nucleus. The scale bar is 1 µm. All the experiments were performed two independent times.

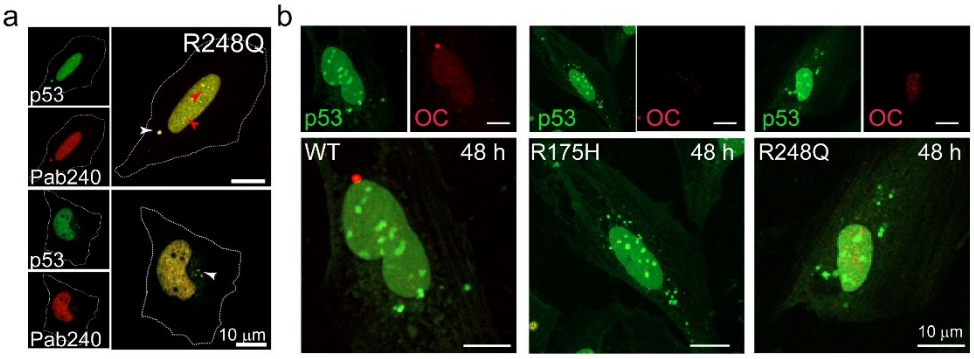


**Figure S6. Characterization of WT and mutant p53 in SaOS2 cells**. **(a)** Immunofluorescence study showing higher colocalization of R248Q GFP p53 condensates with misfolded p53 specific antibody (Pab240). The scale bar is 10 µm. (**b**) Immunofluorescence study showing GFP-p53 WT, GFP-p53 R175H and GFP-p53 R248Q condensates did not show any colocalization with OC antibody (specific to amyloid) in the cytoplasm or nucleus. The scale bar is 10 µm. All the experiments were repeated two times independently.

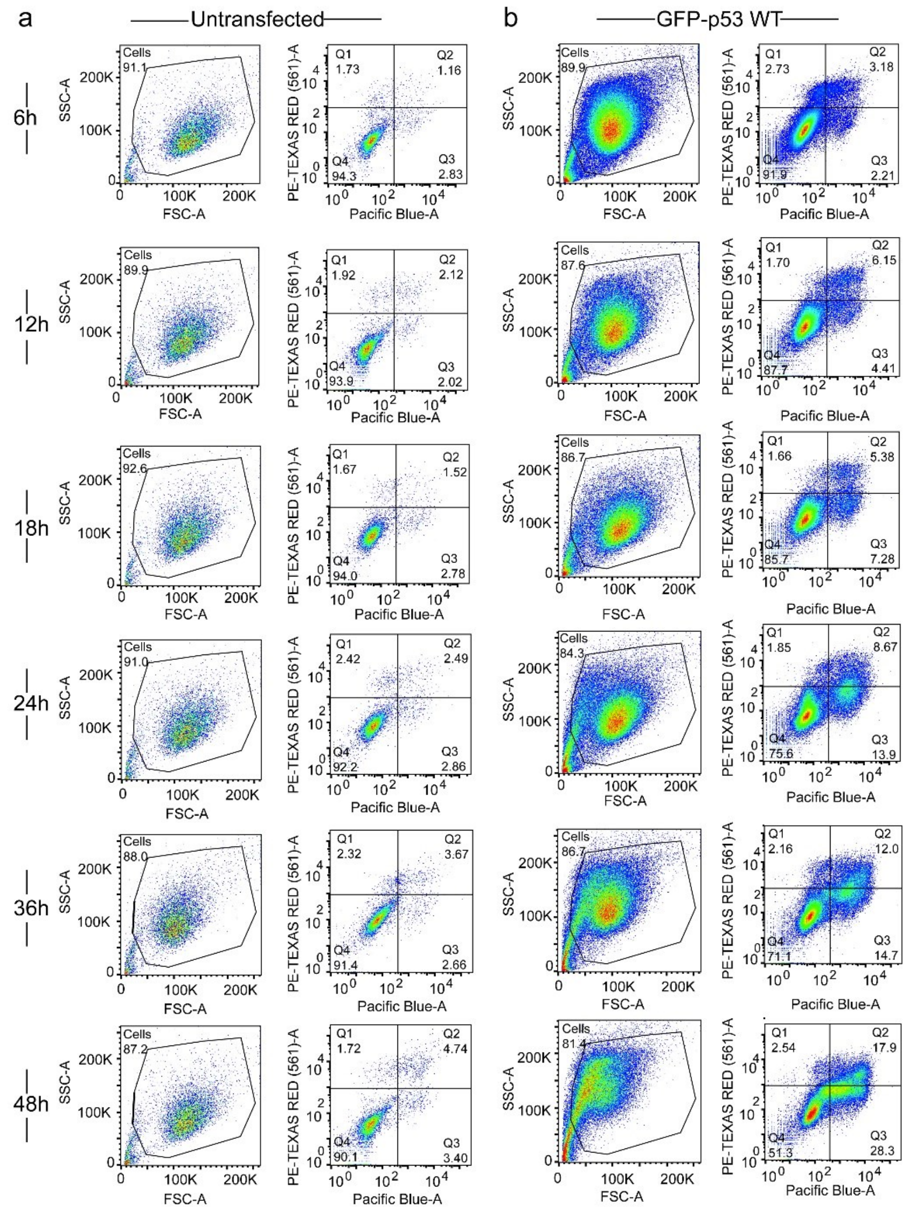


**Figure S7. Time-dependent apoptosis assay for SaOS2 cells expressing WT p53**. **(a-b)** V450 Annexin V-PI assay followed by FACS analysis showing the apoptotic population of cells transfected with GFP-p53 WT over time. Untransfected cells were used as control and the gating was done with respect to the untransfected cell population. GFP-p53 WT positive cells (~$10^4$) were counted to evaluate the apoptotic population. The WT p53 cells showed a lesser apoptotic population at the early time point (6 h, 12 h,18 h), which gradually increased over time (24 h, 36 h, 48 h), indicative of p53 apoptotic function ***(right panel)***. The values are normalized using untransfected control cells ***(left panel)***. The experiment was performed two independent times.

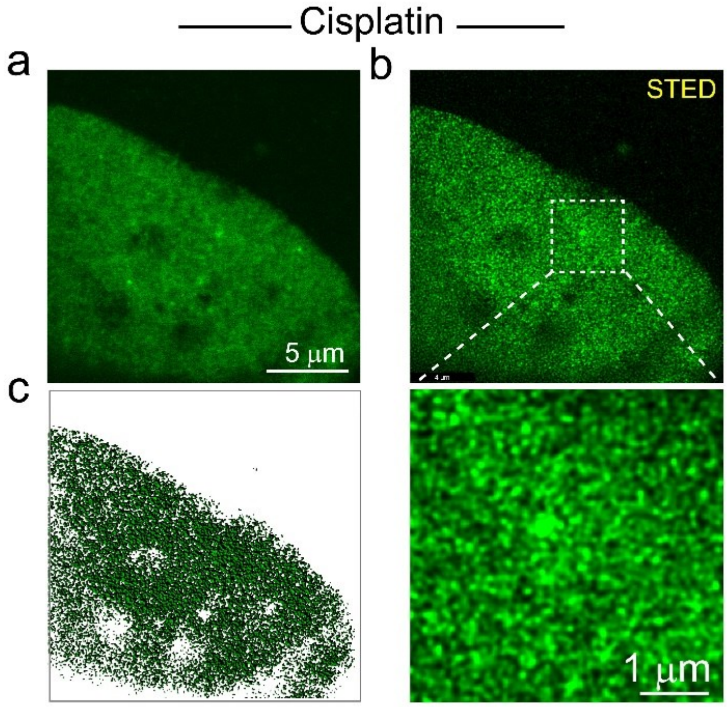


**Figure S8. Super-resolution image of nuclear p53 after stabilization and activation**. **(a)** Confocal images of GFP-p53 in the nucleus of cells treated with Cisplatin. The scale bar is 5 µm. **(b)** Super-resolution microscopy images using STED showing a dense population of p53 condensates in the nucleus. The magnified image of the nuclear condensates is shown in the lower panel. The scale bar is 1 µm. **(c)** The rendered image of p53 nuclear condensates was obtained from STED microscopy images using IMARIS 8.4. All the experiments were repeated two independent times with similar observations.

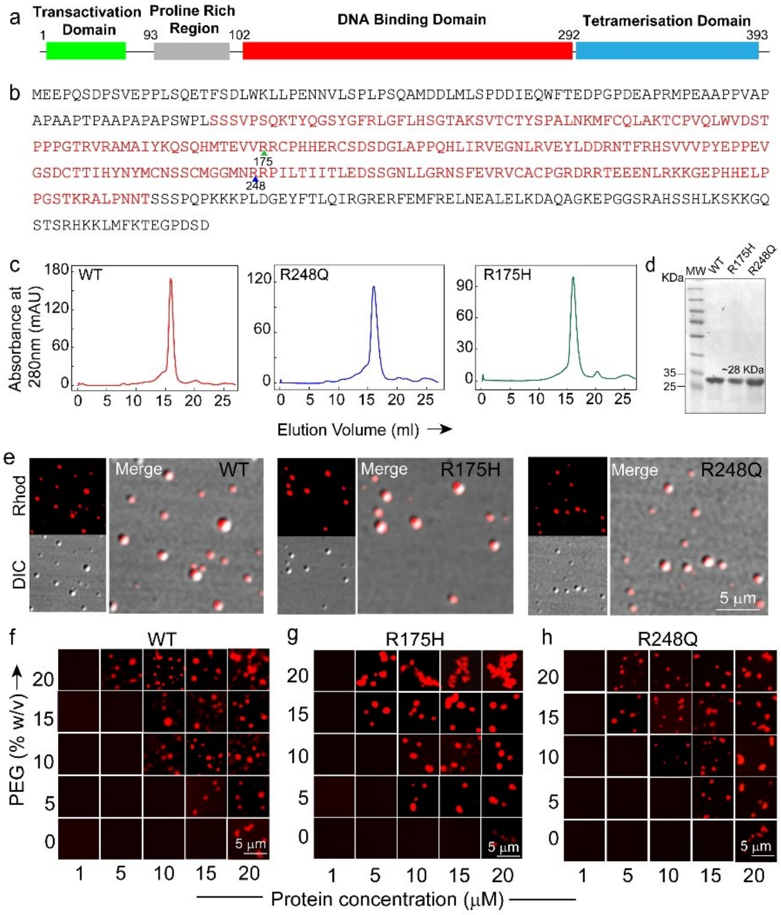


**Figure S9. p53C condensate formation *in vitro*.** **(a)** Schematic representation of the domain organization of full-length p53 showing its organization into N-terminal transactivation domain, Proline-rich region, DNA binding domain and tetramerization domain at the C-terminus. **(b)** The primary sequence of full-length p53 comprises 393 amino acids. The DNA binding region (94-312) (p53C) (red) comprises the two mutant sites R175 (green arrow) and R248 (blue arrow), which are two of the hotspot mutations of p53 associated with cancer. **(c)** Size exclusion chromatography profile of purified p53C of WT, R248Q and R175H after expression in BL21 (DE3) cells, purified via Ni-NTA column chromatography. **(d)** 15% SDS-PAGE of all purified p53C proteins showing the single band at ~ 28 kDa, confirming the purity of the protein. **(e)** Representative confocal microscopy images of 10 µM p53C WT (10% v/v labelled to unlabelled protein), and its hotspot mutants R175H and R248Q, showing condensates formation in the presence of 10% (w/v) PEG-8000 in 50 mM sodium phosphate buffer (pH 7.4). The scale bar is 5 µm. The experiment was repeated three times with similar observations. **(f-h)** The phase regime of the NHS-Rhodamine labelled WT, R175H and R248Q p53C (10% v/v labelled to unlabelled protein) showing the LLPS behaviour at different concentrations of protein in the presence of varying percentages of PEG-8000 (0%, 5%, 10%, 15%, and 20% w/v) in 50 mM sodium phosphate buffer (pH 7.4). The data shows a similar protein concentration requirement for condensate formation for both WT and mutant proteins. The scale bar is 5 µm. The experiment was repeated three independent times with similar observations.

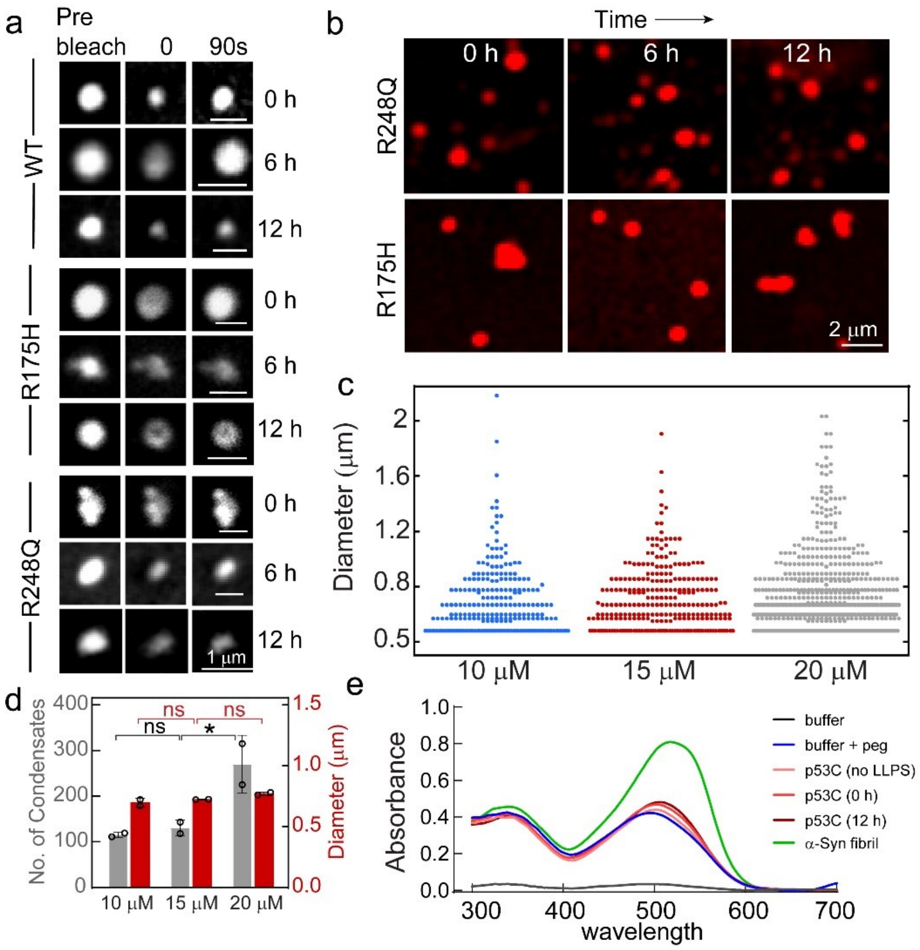


**Figure S10. Characterization of WT and mutant p53C. (a)** Representative confocal images of WT, R175H and R248Q p53C protein condensates during FRAP analysis (pre-bleach, bleach (0 s), and post-bleach (90 s)] at 0 h, 6 h and 12 h after LLPS. The scale bar is 1 µm. The experiment was performed two independent times. **(b)** Representative confocal images of NHS-Rhodamine labelled (10% v/v labelled to unlabeled protein) of 10 µM R175H and R248Q condensates in the presence of 10% (w/v) PEG-8000 at different time intervals. The scale bar is 2 µm. The experiment was performed three independent times. **(c)** Dot plot representing the size distribution of p53C condensates at varying protein concentrations showing a similar size of condensates throughout the concentrations. **(d)** Bar plot representing the number and size of p53C condensates at different protein concentrations showing an increase in number but without any significant change in average size (diameter) with an increase in protein concentrations. Statistical significance was calculated with one-way ANOVA using the Student-Newman-Keuls Multiple Comparison post hoc test (*** $p<0.001$, ** $p<0.01$,* $p<0.05$, ns $p>0.05$). **(e)** Congo red (CR) binding assay showing no significant CR binding to WT p53C condensates immediately after formation (0 h) and 12 h after LLPS. p53 without LLPS and α-Syn fibrils were used as control. The experiment was repeated two independent times.

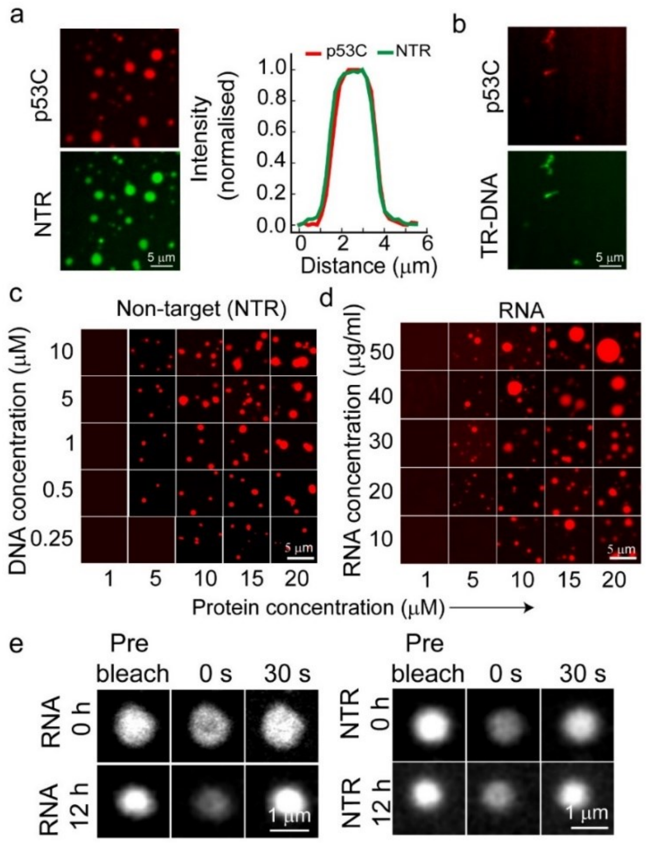


**Figure S11. p53C phase separation in the presence of DNA and RNA**. **(a)** ***(Left panel)*** The confocal microscopy images of 20 μM NHS-Rhodamine labelled [10% (v/v) labelled to unlabelled protein] p53C (red) and Atto-488 labelled [10% (v/v) labelled to unlabelled DNA] NTR-DNA (green) corresponding to multicomponent condensate shown in Fig. 5a (inset). ***(Right panel)*** Line intensity profile showing the colocalization of NHS-Rhodamine labelled p53C with Atto-488 labelled NTR-DNA inside the condensate shown in Fig. 5a (inset). **(b)** The confocal microscopy images showing the remaining small population of foci-like p53 condensates of 20 μM of NHS-Rhodamine labelled [10% (v/v) labelled to unlabelled protein] p53C (red) and Atto-488 labelled [10% (v/v) labelled to unlabelled DNA] TR-DNA (green) corresponding to multicomponent foci shown in Fig. 5a (inset). All the experiments were performed twice with similar observations. **(c, d)** Representative fluorescence microscopy images showing condensate formation by NHS-Rhodamine labelled WT-p53C [10% (v/v) labelled to unlabelled protein] with varying concentration of NTR-DNA **(c)** and Poly U-RNA **(d)** in the presence of 10% (w/v) PEG-8000 in 50 mM sodium phosphate buffer (pH 7.4). The scale bar is 5 μm. **(e)** Representative confocal images of p53C condensates formed in the presence of RNA and NTR-DNA during FRAP analysis (pre-bleach, bleach (0 s), and post-bleach (30 s)] at 0 h and 12 h after LLPS. The scale bar is 1 μm. The experiment was performed two independent times.

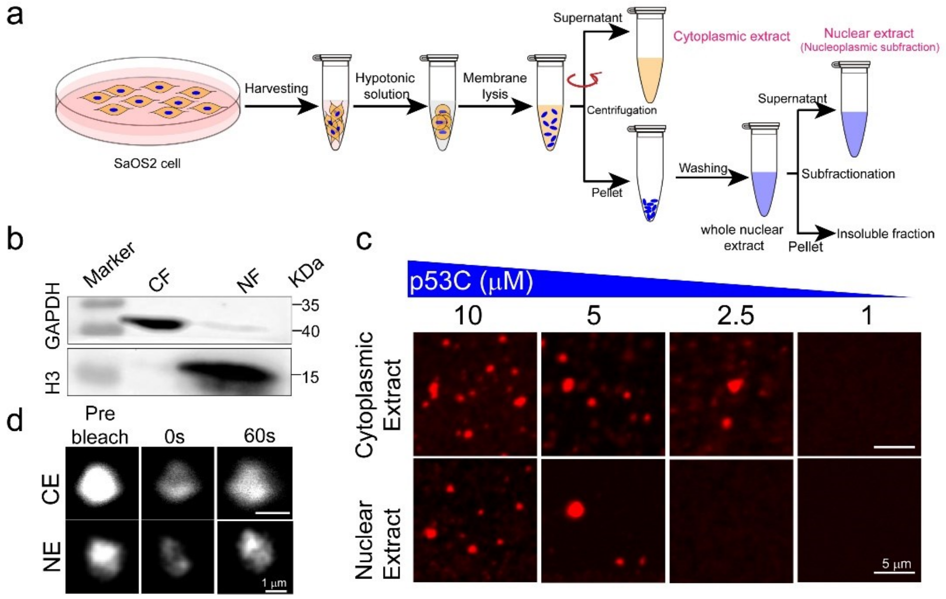


**Figure S12**. **p53C condensate formation in the presence of nuclear and cytoplasmic extracts of SaOS2.** **(a)** The schematic representation showing the separation protocol of nuclear and cytoplasmic fractions from SaOS2 cells. **(b)** Western blot of SaOS2 cells extracts showing cytoplasmic marker GAPDH in the cytoplasmic fraction (CF) and nuclear marker H3 in the nuclear fraction (NF) only, indicating the separation of the cytoplasmic and nuclear fractions. **(c)** Representative fluorescence microscopy images of WT p53C [10% (v/v) NHS-Rhodamine labelled to unlabelled protein] condensates at different protein concentrations in the presence of nuclear (NE) and cytoplasmic extract (CE) of SaOS2 cells. The scale bar is 5 μm. The experiment was repeated three times with similar observations. **(d)** Representative images of p53C condensates formed in the cytoplasmic (***upper***) and nuclear (***lower***) extract at pre-bleach, bleach (0 s) and post-bleach states (60 s). The scale bar is 5 μm. The experiment was performed two independent times.

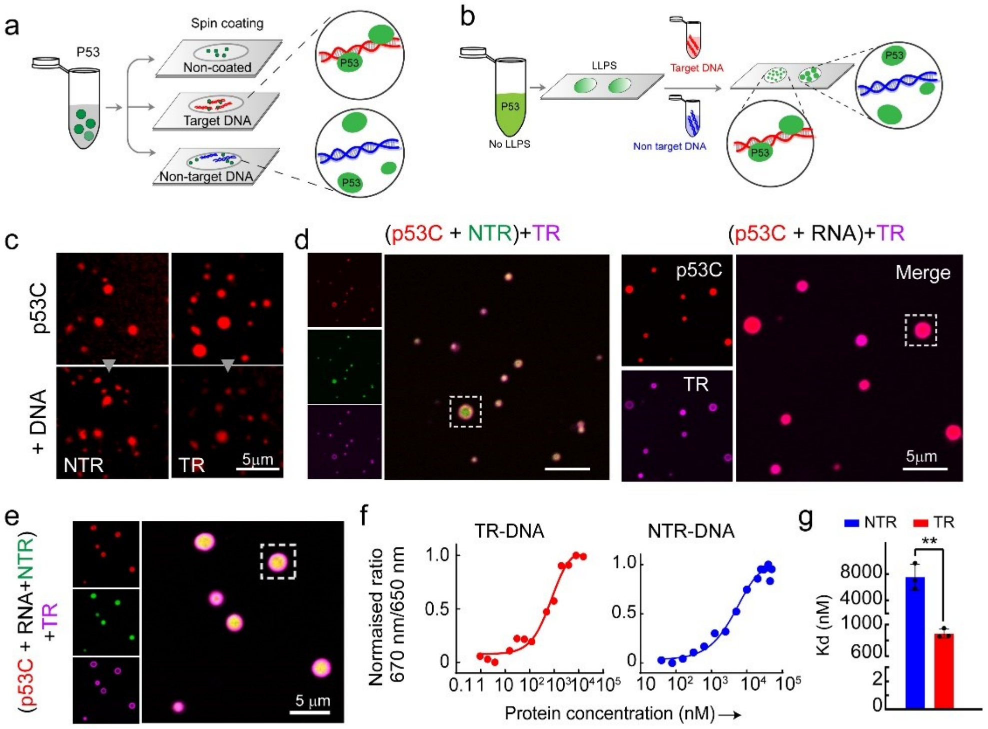


**Figure S13. Effect of TR-DNA on pre-formed p53 condensates. (a-b)** Schematic representations of the experiments performed to examine the effect of target (TR) and non-target DNA (NTR-DNA) on pre-formed p53C condensates. **(a)** Pre-formed LLPS mixture of p53C formed in the presence of 10% (w/v) PEG-8000 was spotted on glass coverslips that were pre-coated with either TR-DNA or NTR-DNA. **(b)** p53C in the presence of 10% (w/v) PEG-8000 was spotted in chambers on coverslips mounted with silicone isolators (Grace Biolabs). After phase separation, TR-DNA and NTR-DNA were added to the solution in separate chambers and observed under the microscope. **(c)** The confocal microscopy images of p53C condensates in the presence of 10% (w/v) PEG-8000 ***(upper panel)*** showing increase in partitioning of p53 into the condensates upon addition of NTR-DNA, while decrease in partitioning upon addition of TR-DNA (***lower panel)***. The scale bar is 5 μm. The experiment was performed twice with similar results. **(d-e)** Representative confocal microscopy images showing p53C condensates which remained after the addition of Atto 647N labelled TR-DNA to the pre-formed p53C-NTR, p53C-RNA and p53C-NTR-RNA condensates. NTR-DNA was labelled with Atto 488. The inset white box shows condensates represented in the main Figure 6 h-j. **(f)** Spectral shift measurement of p53C (10 μM) in the presence of TR-DNA and NTR-DNA showing binding affinity of p53C with nucleic acids. **(g)** Bar graph representing the binding affinity ($K_d$) of p53C with TR-DNA and NTR-DNA showing a significantly higher binding affinity for TR-DNA. Error bars represent mean ± s.e.m for n=3 independent experiments. The statistical significance was estimated with an unpaired t-test with 95% confidence interval (***p< .001, **p<.002).

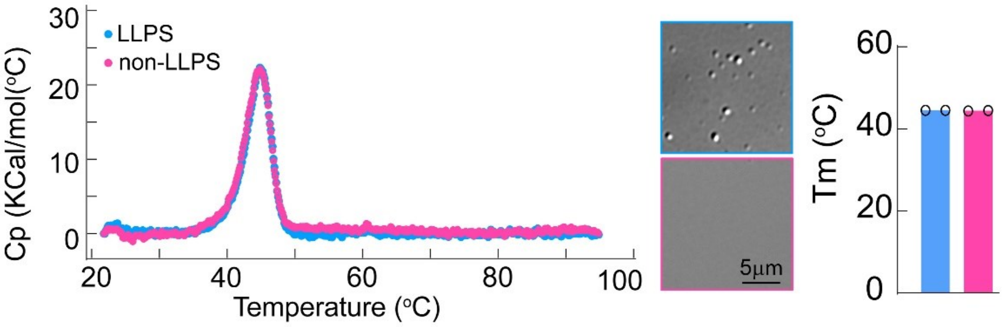


**Figure S14. Differential Scanning calorimetry (DSC) of p53C in LLPS and non-LLPS states.** The DSC data showing no significant change in melting temperature ($T_m$). DIC image showing p53C in LLPS and Non-LLPS state (***middle panel***) and corresponding melting $T_m$ (***right panel***).

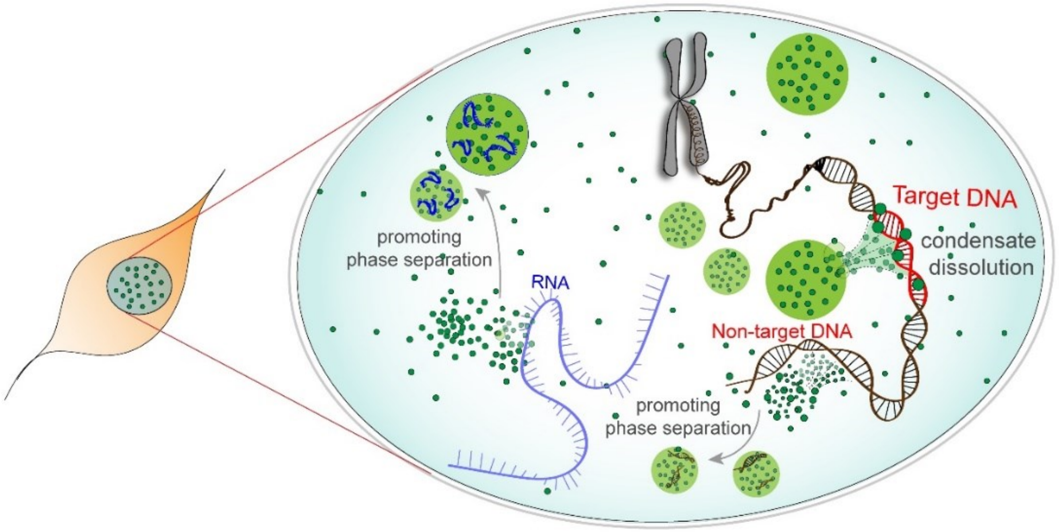


**Figure S15. Nuclear model of p53 condensates**. Schematic representation of the possible mode of operation of p53 in the nuclear microenvironment, where p53 formed condensates, which is further facilitated by the interaction with non-target DNA and RNA. However, the interaction/vicinity with target DNA, p53 condensates get dissolved and bind to the target DNA for the activation of downstream regulatory functions.

**Table S1. Primers used for site-directed mutagenesis**

| Name | Sequence Details | Source |
|---|---|---|
| R175H- Forward | 5'- GGAGGTTGTGAGGCATTGCCCCC - 3' | Sigma |
| R175H- Reverse | 5'- GGGGGCAATGCCTCACAACCTCC - 5' | Sigma |
| R248Q-Forward | 5'- GGGCGGCATGAACCAGAGGCCCATCCTCAC - 3' | Sigma |
| R248Q-Reverse | 5'- GTGAGGATGGGCCTCTGGTTCATGCCGCCC - 3' | Sigma |

**Table S2. Sequences of Target, Non-Target DNA and RNA used**

| Nucleic Acid | Sequence details | | Source |
|---|---|---|---|
| Target DNA | Forward | 5'- ATCAGGAACATGTCCCAACATGTTGAGCTC - 3' | Sigma |
| | Reverse | 5’ - GAGCTCAACATGTTGGGACATGTTCCTGAT - 3’ | Sigma |
| Non-Target DNA | Forward | 5’ - AATATGGTTTGAATAAAGAGTAAAGATTTG - 3’ | Sigma |
| | Reverse | 5' - CAAATCTTTACTCTTTATTCAAACCATATT - 3' | Sigma |
| RNA | Poly U RNA | | Sigma#9528 |

**Table S3. Primers for qRT-pCR genes (GAPDH and P21)**

| Gene | REVERSE (5'-3') | FORWARD (5'-3') | Source |
|---|---|---|---|
| GAPDH | GGGTTCGAAATGAGGATG | GGGTTCGAAATGAGGATG | Sigma |
| P21 | GGTAGAAATCTGTCATGCTG | AAGACCATGTGGACCTGT | Sigma |

**SUPPLEMENTARY MOVIE LEGENDS:**

**Supplementary Movie 1**: Time-lapse video of WT GFP-p53 transfected HeLa cells showing cytoplasmic and nuclear condensates exhibiting dynamic nature at 18 h. Time is represented by hh:mm:ss.

**Supplementary Movie 2:** Time-lapse video showing fusion of cytoplasmic condensates in HeLa cells transfected with WT GFP-p53. Time is represented by hh:mm:ss.

**Supplementary Movie 3**: Time-lapse video showing fusion of cytoplasmic condensates in SaOS2 cells transfected with WT GFP-p53. Time is represented by hh:mm:ss.

**Supplementary Movie 4:** Time-lapse video of a single SaOS2 cell, exhibiting spatiotemporal localization of p53, showing the formation of cytoplasmic condensates followed by cell death. Scale bar is 10 μm. Time is represented by hh:mm:ss.

**Supplementary Movie 5:** Lattice light-sheet microscopy imaging showing the dynamic nature of nuclear condensates in SaOS2 cells transfected with WT GFP-p53 and 18 h and 48 h. Time is represented by mm:ss.

**Supplementary Movie 6:** Time-lapse video showing the formation of p53C condensates at 20 μM (NHS-Rhodamine labelled p53C 1:10::labelled:unlabelled) *in vitro* with time. Time is represented by mm:ss. Scale bar represents 5 μm.

#### References


1 Erdős, G. & Dosztányi, Z. Analyzing protein disorder with IUPred2A. *Current Protocols in Bioinformatics* **70**, e99 (2020).

2 Mészáros, B., Erdős, G. & Dosztányi, Z. IUPred2A: context-dependent prediction of protein disorder as a function of redox state and protein binding. *Nucleic Acids Research* **46**, W329-W337 (2018).

3 Miskei, M., Horvath, A., Vendruscolo, M. & Fuxreiter, M. Sequence-based prediction of fuzzy protein interactions. *Journal of molecular biology* **432**, 2289-2303 (2020).

4 Letunic, I. & Bork, P. 20 years of the SMART protein domain annotation resource. *Nucleic acids research* **46**, D493-D496 (2018).

5 Jumper, J. *et al.* Highly accurate protein structure prediction with AlphaFold. *Nature* **596**, 583-589 (2021).

6 Wang, Y., Rosengarth, A. & Luecke, H. Structure of the human p53 core domain in the absence of DNA. *Acta Crystallographica Section D: Biological Crystallography* **63**, 276-281 (2007).

7 DeLano, W. L. Pymol: An open-source molecular graphics tool. *CCP4 Newsletter on Protein Crystallography* **40**, 82-92 (2002).

8 Lowe, D. G. Distinctive Image Features from Scale-Invariant Keypoints. *International Journal of Computer Vision* **60**, 91-110 (2004).

9 Ghosh, S. *et al.* p53 amyloid formation leading to its loss of function: implications in cancer pathogenesis. *Cell Death & Differentiation* **24**, 1784-1798 (2017).

10 Ray, S. *et al.* α-Synuclein aggregation nucleates through liquid–liquid phase separation. *Nature chemistry* **12**, 705-716 (2020).

11 Blom, H. & Widengren, J. Stimulated emission depletion microscopy. *Chemical reviews* **117**, 7377-7427 (2017).

12 Hell, S. W. & Wichmann, J. Breaking the diffraction resolution limit by stimulated emission: stimulated-emission-depletion fluorescence microscopy. *Optics Letters* **19**, 780-782 (1994).

13 Schoonderwoert, V., Dijkstra, R., Luckinavicius, G., Kobler, O. & van der Voort, H. Huygens STED Deconvolution Increases Signal-to-Noise and Image Resolution towards 22 nm. *Microscopy Today* **21**, 38-44 (2013).

14 Jui-Cheng, Y., Fu-Juay, C. & Shyang, C. A new criterion for automatic multilevel thresholding. *IEEE Transactions on Image Processing* **4**, 370-378 (1995).

15 Sezgin, M. & Sankur, B. l. Survey over image thresholding techniques and quantitative performance evaluation. *Journal of Electronic imaging* **13**, 146-168 (2004).

16 Sauvola, J. & Pietikäinen, M. Adaptive document image binarization. *Pattern Recognition* **33**, 225-236 (2000).

17 Neerad, P., Sumit, M., Ashish, S. & Madhuri, J. Adaptive local thresholding for detection of nuclei in diversity stained cytology images. *2011 International Conference on Communications and Signal Processing.* 218-220 (2011).

18 Arganda-Carreras, I. *et al.* Trainable Weka Segmentation: a machine learning tool for microscopy pixel classification. *Bioinformatics* **33**, 2424-2426 (2017).

19 Sengupta, S. *et al.* p53 amyloid pathology is correlated with higher cancer grade irrespective of the mutant or wild-type form. *Journal of Cell Science* **136,** 17 (2023).

20 Milkovic, N. M. & Mittag, T. Determination of protein phase diagrams by centrifugation. *Intrinsically Disordered Proteins: Methods and Protocols* **2141**, 685-702 (2020).

21 Langer, A. *et al.* A New Spectral Shift-Based Method to Characterize Molecular Interactions. *ASSAY and Drug Development Technologies* **20**, 83-94 (2022).

22 Senichkin, V. V., Prokhorova, E. A., Zhivotovsky, B. & Kopeina, G. S. Simple and efficient protocol for subcellular fractionation of normal and apoptotic cells. *Cells* **10**, 852 (2021).

23 Freibaum, B. D., Messing, J., Yang, P., Kim, H. J. & Taylor, J. P. High-fidelity reconstitution of stress granules and nucleoli in mammalian cellular lysate. *Journal of Cell Biology* **220**, e202009079 (2021).
